# Where were the Caribbean mangroves during the Last Glacial Maximum? A microtopographical approach

**DOI:** 10.1101/2025.01.13.632686

**Authors:** Valentí Rull

## Abstract

During the Last Glacial Maximum (LGM), when global sea levels dropped by ∼132 m, the Caribbean continental shelf was fully exposed, which drastically reduced the flat topographical habitat necessary for mangrove growth. It has been proposed that mangroves survived in flat microsites beyond the shelf break and later expanded from these microrefugia to their current distribution after the LGM. However, this hypothesis remains untested. This study aims to identify potential refugia by locating flat areas around the -132 m isobath using Global Multiresolution Topography (GMRT) images. A significant ∼200-km-long potential refugium was identified on the northern Trinidad (NT) shelf, along with several scattered kilometer-scale microrefugia near the Cariaco Basin (CB) in northeastern Venezuela. Additionally, two isolated prospective microrefugia were detected in northern Colombia (NC) and western Hispaniola (WH). Bioclimatically, all these potential LGM refugia were suitable for mangrove growth. The remaining LGM Caribbean coasts were considered unsuitable for mangrove growth. The NT refugium, along with the CB microrefugia, may have served as the primary sources for subsequent mangrove expansion. This expansion was likely facilitated by postglacial sea-level rise and the SE-NE Caribbean Current (CC), which would have acted as a major agent for propagule dispersal. This microtopographical survey not only supports the microrefugial hypothesis but also narrows the focus to the most promising areas, significantly reducing the time, effort and resources required for future seismic and coring campaigns.

## 1. Introduction

Mangroves are tropical and subtropical intertidal forested ecosystems that are vital for maintaining coastal biodiversity, providing a range of ecological services, and serving as the most significant blue-carbon ecosystems, contributing to the mitigation of global warming. These ecosystems are among the most endangered on Earth (Worthington et al., 2020). Recent assessments indicate that global mangrove coverage has decreased by 3.4% in less than 25 years (1996–2020) due to both natural and human-induced deforestation (Bunting et al., 2022). If these depletion rates continue, mangroves face a significant risk of substantial decline during this century, posing a serious threat to their long-term survival (Duke et al., 2017).

Mangroves thrive around the average sea level, forming vegetation gradients that range from coastal marine habitats to the upper limits of normal tides in continental environments. They are especially well-developed in extensive, flat sedimentary areas, such as deltas or carbonate platforms, where wave action is weak or absent (Thom, 1984; Schaeffer-Novelli et al., 2002; Mazda et al., 2007). In contrast, mangroves do not develop in non-flooded or topographically complex environments, such as steep coastal slopes, rocky substrates, or littoral cliffs. A recent review by Ellison et al. (2024) reports that mangroves worldwide occupy elevation ranges between -0.90 and 3.60 m (−0.12 and 1.25 m on average) and extend inland from a few hundred meters to ∼15 km (average 3.26 km). According to these data, the mean slope of flat areas (FAs) supporting mangroves ranges between 0.01% and 0.45% (average 0.12%), representing 35% to 267% (average 78%) of the total tidal range.

In the Caribbean region (Fig. 1), mangroves cover a total area of approximately 14,700 km^2^, accounting for ∼10% of the world’s mangrove surface (Bunting et al., 2022). They are distributed along all Caribbean coasts, with the largest extents found in Cuba, Venezuela, Colombia, and Panama (1,500–3,600 km^2^). In contrast, all other countries have mangrove areas of 600 km^2^ or less, with 15 countries having less than 100 km^2^. The primary mangrove-forming trees in the region include the red mangrove (*Rhizophora*, Rhizophoraceae), black mangrove (*Avicennia*, Acanthaceae), and white mangrove (*Laguncularia*, Combretaceae). Additional components, such as the tea mangrove (*Pelliciera*, Tetrameristaceae) and the mangrove fern (*Acrostichum*, Pteridaceae), are also typical but occupy marginal environments.

**Figure 1.**
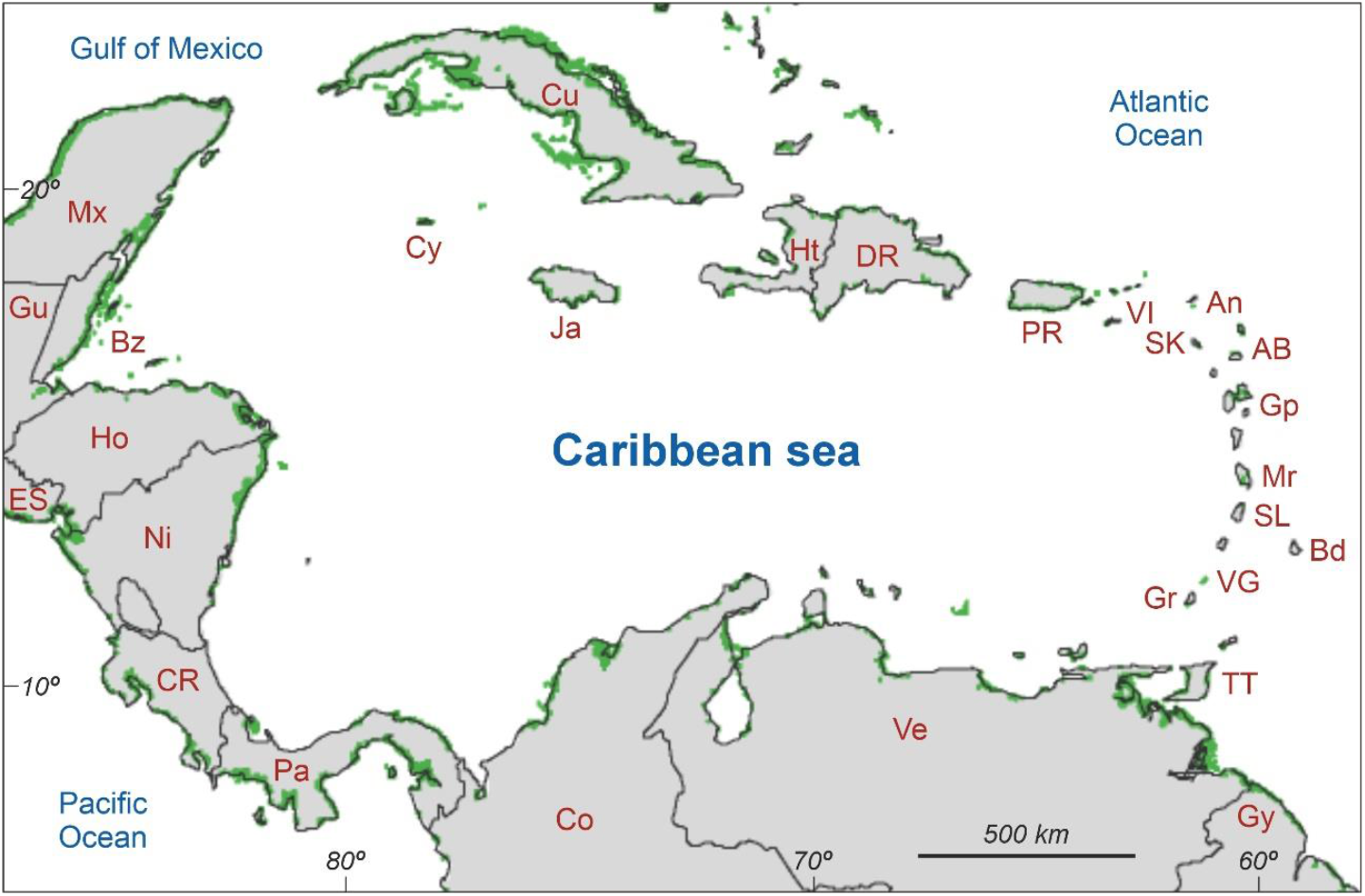
Map of the Caribbean mangroves (green patches), according to Bunting et al. (2022). Downloaded from NASA Landsat 5-TM (https://earthobservatory.nasa.gov/images/47427/mapping-mangroves-by-satellite). Countries/islands: AB, Antigua & Barbuda; An, Anguilla (UK); Bd, Barbados; Bz, Belize; Co, Colombia; CR, Costa Rica; Cu, Cuba; Cy, Cayman Islands (UK); DR, Dominican Republic; ES, El Salvador; Gp, Guadeloupe (France); Gu, Guatemala; Gy, Guyana; Ht, Haiti; Ja, Jamaica; Mr., Martinique (France); Mx, Mexico; Ni, Nicaragua; Pa, Panama; PR, Puerto Rico; SK, Saint Kitts & Nevis; SL, Saint Lucia; TT, Trinidad & Tobago; Ve, Venezuela; VG, Saint Vincent & The Grenadines; VI, Virgin Islands (USA/UK)

The Caribbean mangroves originated during the Middle Eocene (50–40 Ma) and were initially dominated by *Pelliciera* until the Eocene-Oligocene transition (EOT), when *Rhizophora* became the primary mangrove-forming tree. The Neogene marked a period of diversification, culminating in the current mangrove composition during the Pliocene. Throughout this time, mangrove evolution and biogeography were heavily influenced by regional tectonics, including the movement of the Caribbean plate, continental drift, and climatic/eustatic changes, such as the EOT cooling and associated sea-level fall. In the Quaternary, external drivers included the Pleistocene glacial/interglacial cycles and human activities, which began to have a regional impact around 6–5 ka BP. The detailed evolutionary history of mangroves can be found in Rull (2024a).

Pleistocene paleoecological records of in situ Caribbean mangroves are rare. The oldest such record, found on an exposed Jamaican terrace, consists of mangrove root scars dated to ∼130 ka BP (MIS 5e highstand), a period when sea levels were several meters higher than today (Mitchell et al., 2001). Another record, from the Cariaco Basin (Fig. 2), dates to the Last Glaciation (approximately 68–28 ka BP), encompassing Heinrich Events H6 to H3 (González et al., 2008; González & Dupont, 2009). This record, obtained from a depth of 900 m, shows pollen of *Rhizophora* and *Acrostichum* consistently contributing less than 4% of the total pollen assemblage. Notably, there is a gap in the records that includes the Last Glacial Maximum (LGM; 27–19 ka BP; Clark et al., 2009). The next in situ record of mangroves, dominated by *Rhizophora* and *Avicennia*, corresponds to the earliest Holocene (12.4–11.8 ka BP) in the Orinoco Delta (Pocknall & Jarzen, 2024). As such, the LGM remains absent from Caribbean mangrove records.

**Figure 2.**
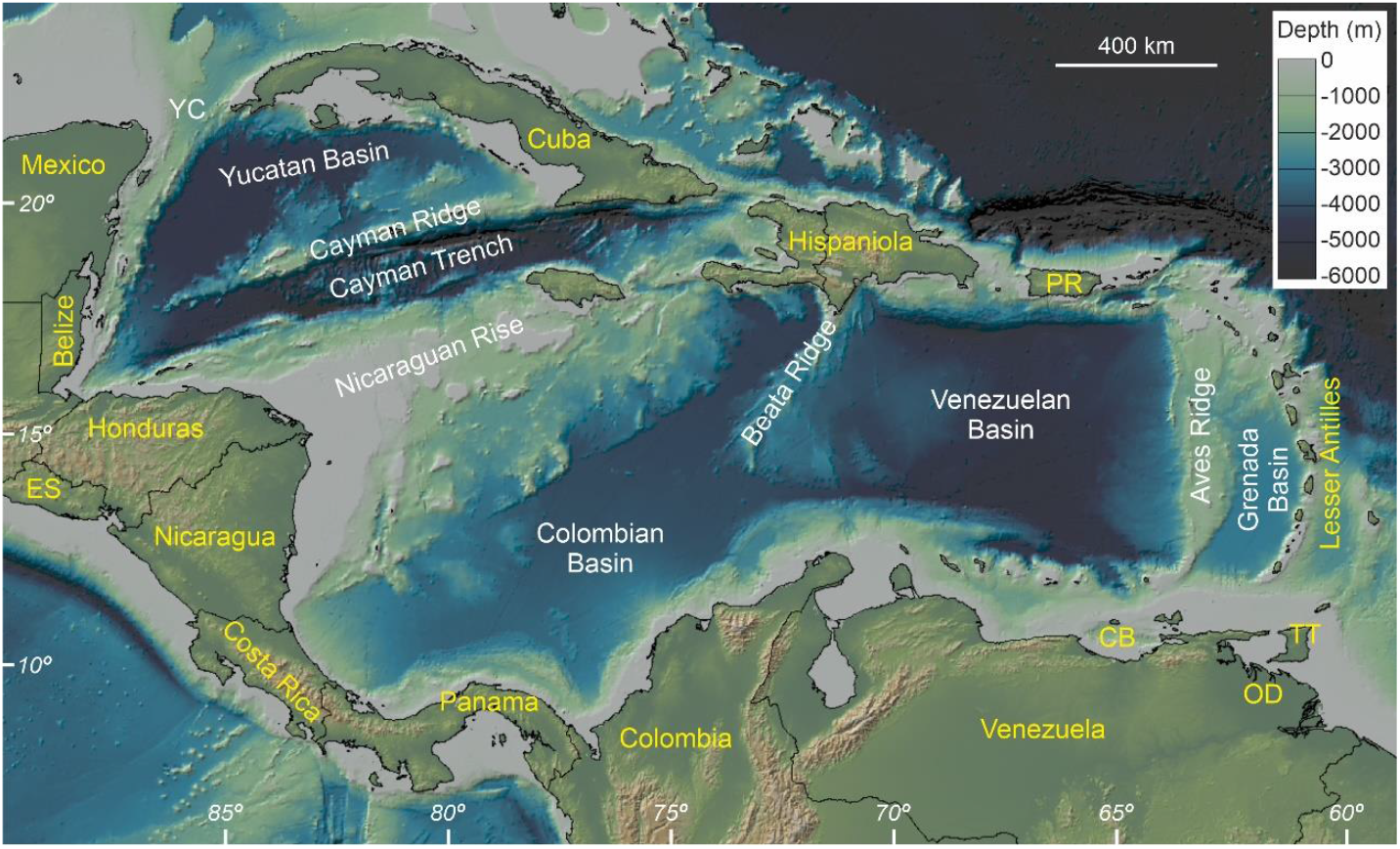
Reference map for this study. Overview of the Caribbean seafloor using Global Multi-Resolution Topography (GMRT) (see methods). Abbreviations: CB, Cariaco Basin; ES, El Salvador; Ja, Jamaica; OD, Orinoco Delta; PR, Puerto Rico; TT, Trinidad & Tobago; YC, Yucatan Channel.

During the LGM, global relative sea levels (RSL) were 130–134 m below present levels (Lambeck et al., 2014; Spratt & Liesecki, 2016). This substantial drop exposed large portions of continental shelves worldwide, which typically reach maximum depths of ∼200 m (IHO, 2019). The exposure of these shelves likely had a significant impact on mangrove distribution, as most mangroves today grow on flat upper shelf environments, while LGM coastlines were located near or on steeper continental slopes, which are topographically unsuitable for mangroves. In the Caribbean, it has been hypothesized that LGM mangroves persisted in microrefugia beyond the shelf break in areas with favorable microenvironmental conditions. From these refugia, mangroves are thought to have expanded along Caribbean coasts during the postglacial sea-level rise, ultimately reaching their current distribution (Rull, 2024b). However, this hypothesis remains untested due to a lack of suitable paleoecological records.

To identify these potential LGM mangrove refugia, a high-resolution bathymetric survey is required to locate suitable FAs around the LGM-RSL. This study aims to identify these FAs around -132 m (the midpoint of the -130 to -134 m range) using a microtopographical bathymetric approach. This survey can help set the bases for understanding the biogeographical patterns of LGM mangroves and their subsequent range shifts during the Late Glacial and Holocene, leading to their present-day distribution. In addition, it is able to identify the best prospects for further coring campaigns aimed at locating the LGM mangrove communities using paleoecological methods, notably pollen analysis.

## 2. Study site and methods

### 2.1. Overview of the Caribbean seafloor

The Caribbean seafloor is a deep and complex surface shaped by the tectonic dynamics of the Caribbean microplate, located between the North and South American plates and the Pacific Cocos and Nazca plates (Mann, 2021). It is subdivided into four major basins, arranged from west to east: the Yucatan Basin (∼4,350 m maximum depth), the Colombian Basin (∼4,160 m), the Venezuelan Basin (∼5,630 m), and the Grenada Basin (∼3,000 m) (Fig. 2). These basins are separated by three significant submarine elevations: the Nicaraguan Rise, the Beata Ridge, and the Aves Ridge. The deepest point in the Caribbean is the Cayman Trench (∼7,690 m), part of the Cayman Basin, which is separated from the Yucatan Basin by the Cayman Ridge.

The most extensive continental shelves are found along the western and southern coasts, particularly in the regions of Honduras/Nicaragua and the northwestern and northeastern coasts of Venezuela, including Trinidad and Tobago. The least extensive continental shelves are along the Mexican Caribbean coast, as well as the coasts of Costa Rica, Panama, northern Colombia, and north-central Venezuela. Among the islands, Cuba, Puerto Rico, and the northernmost and southernmost Lesser Antilles have notable continental shelf extents relative to their sizes. In contrast, for most other islands, the shelf break is located close to their coasts or just a few kilometers offshore. There is a strong correlation between the extent of continental shelves and the occurrence of mangroves, as evidenced by a comparison of Figures 1 and 2.

### 2.2. Bathymetric image analysis

This survey focuses on identifying potential FAs for mangrove growth during the Last Glacial Maximum (LGM) lowstand using high-resolution bathymetric data. The primary tool for this analysis is the Global Multi-Resolution Topography (GMRT) dataset (Ryan et al., 2009), version 4.3.0, accessed through GeoMapApp 3.7.4 (www.geomapapp.org). This platform is hosted by the Lamont-Doherty Earth Observatory of Columbia University (USA) under a Creative Commons Attribution Noncommercial Share Alike 3.0 license. The analysis considers both horizontal (bathymetric contours) and vertical (land-sea elevation profiles) components and is conducted at three resolutions: macrotopographic, mesotopographic, and microtopographic.

The macrotopographic analysis utilizes the -132 m contour to delineate the portion of the continental shelf exposed during the LGM. Meso- and microtopographic analyses focus on a meter-scale depth range around this contour to identify FAs more suitable for mangrove growth. The depth range for meso- and microtopographic analyses was determined based on tidal amplitude.

The Caribbean is generally considered a microtidal region, with a tidal range below 20 cm (Kjerfve, 1981; Torres & Tsimplis, 2011). However, local variations exist, with maximum amplitudes of up to 2.6 m recorded along the Orinoco Delta coasts (Warne et al., 2002). During the Holocene, models suggest that Caribbean tidal ranges remained relatively stable, except around 9 ka BP, when tidal amplitudes reached 1–1.5 m along the western coasts near the Yucatan and Colombian basins (Hill et al., 2011; Khan et al., 2017). For the LGM, some modeling studies propose that tidal amplitudes in the Caribbean were similar to those of today (Thomas & Sündermann, 1999), while others estimate increases of 0.5–1 m along the Central American coasts (Egbert et al., 2004).

These dimensions align with the elevation ranges for mangrove gradients reported globally by Ellison et al. (2024), which extend up to 4.5 m, with average slopes of 0.12% (0.01–0.45%). Of this range, 20% or less lies below mean sea level. For an average LGM-RSL of -132 m, this corresponds to a gradient from approximately -128 m (upper mangrove limit; UML) to -133 m (lower mangrove limit; LML) over an average distance of 4.17 km (minimum 1.11 km). These conditions maximize the likelihood of identifying larger FAs suitable for mangrove growth while minimizing the chance of overlooking smaller but favorable FAs.

In the first step of this analysis, prospective FAs were identified as areas where the meter-scale isobaths from -128 to -133 m were more widely spaced. For this mesotopographic analysis, the Caribbean region between 8° and 23° N latitude and 60° to 90° W longitude was divided into 30 zones measuring 3° x 5° each (see Supplementary Material). Steep coasts and cliffs were identified as thin lines, reflecting tightly clustered isobaths along the same paths. In contrast, prospective FAs appeared as thick lines or dark spots, indicating wider isobath spacing over larger surfaces. In the second step, the microtopographic analysis, the identified prospective areas were magnified and examined in detail for the -128 to -133 m interval. This analysis used both isobaths and land-sea profiles and was compared against the elevation gradients reported by Ellison et al. (2024).

## 3. Results

### 3.1. Macrotopography

During the LGM, the shallow (<100 m) and discontinuous Caribbean continental shelf was mostly exposed (Fig. 3). The greatest exposure occurred in southern Cuba, the Nicaraguan Rise, and the northwestern and northeastern coasts of Venezuela, including Trinidad. In contrast, areas where the continental shelf is absent or where the shelf break lies close to the coast—such as Mexico/Belize, the Panama Isthmus, north-central Venezuela, and Hispaniola (Haiti and the Dominican Republic)—experienced minimal exposure. The Nicaraguan Rise is the only submarine elevation with exposed areas located far from present-day coasts.

**Figure 3.**
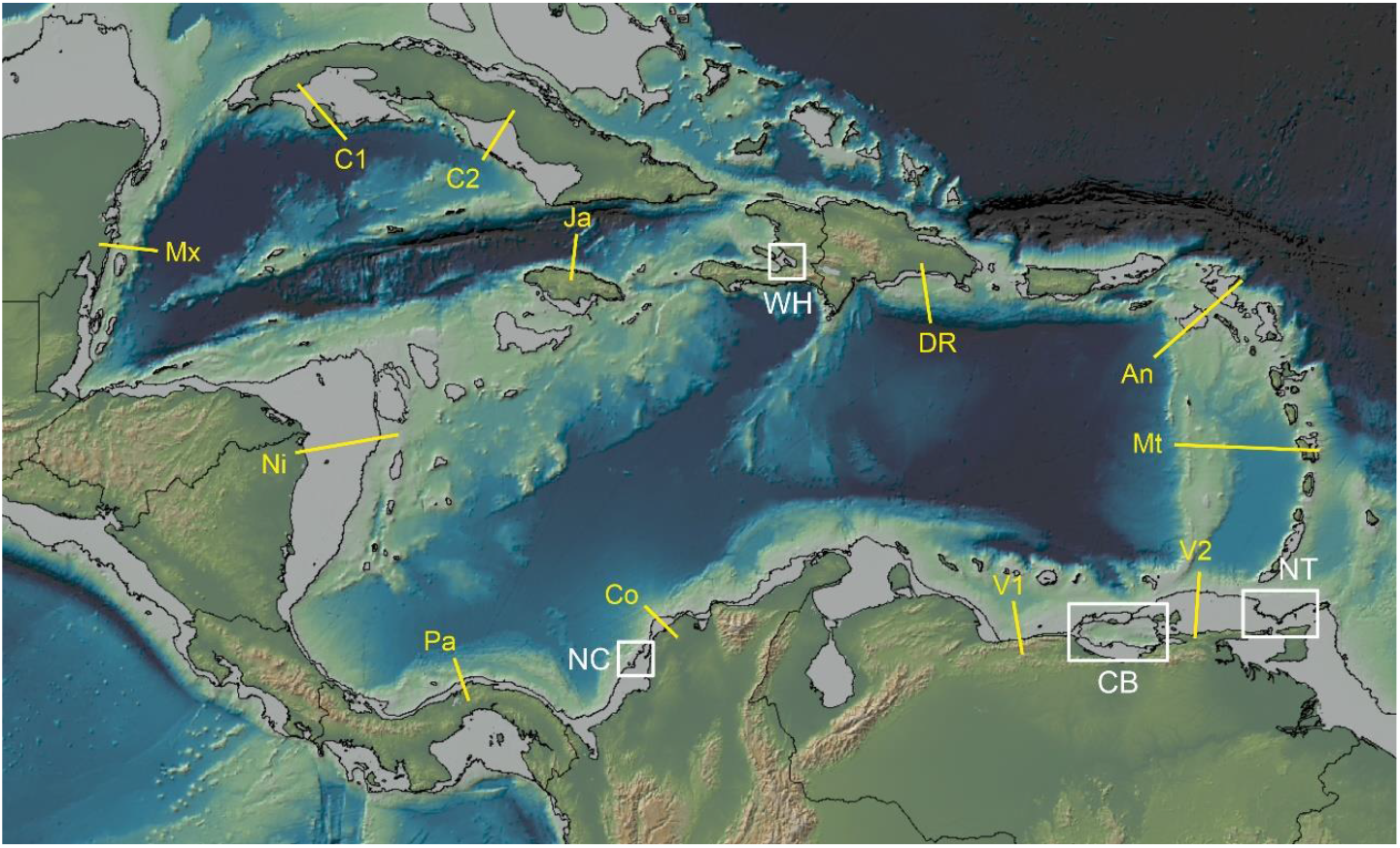
Map showing the -132 m contour (black line beyond the current coasts), the location of the land-sea elevation profiles displayed in Fig 4 (yellow lines) and the most prospective areas resulting from mesotorpographical analysis (white boxes), which are magnified in Figs. 5-7. Base map as in Fig. 2. Profiles: An, Anguilla; Co, Colombia; C1 and C2, Cuba; DR, Dominican Republic; Ja, Jamaica; Mt, Martinique; Mx, Mexico; Ni, Nigaragua; Pa, Panama; V1 to V3, Venezuela. Prospective areas: CB, Caariaco Basin; NC, northern Colombia; NT, northern Trinidad; WC, western Hispaniola.

A set of land-sea elevation profiles was selected around the Caribbean coasts to better visualize the submarine macrotopography of the continental shelf and slope (Fig. 4). The exposed shelf extends more than 100 km offshore in areas such as Cuba, northeastern Venezuela/Trinidad, and Nicaragua, where the shelf break is abrupt, the continental slope is steep, and the transition is continuous. In contrast, regions like Mexico and the Dominican Republic have a narrower exposed shelf, typically only a few tens of kilometers wide. The slope in these areas is composite, featuring two major steps separated by an intermediate plateau between approximately 1,200 and 1,800 m. In regions with high coastal ranges exceeding 2,000 m, such as north-central Venezuela, there is no significant shelf, and the continental slope directly continues from the mountain slope. The most complex profiles are found along the easternmost Caribbean coasts, where the shelf is fragmented in the Lesser Antilles, and the abyssal zone is interrupted by the Aves Ridge, as seen near Anguilla and Martinique. In all cases, the LGM-RSL was located offshore, beyond the shelf break.

**Figure 4.**
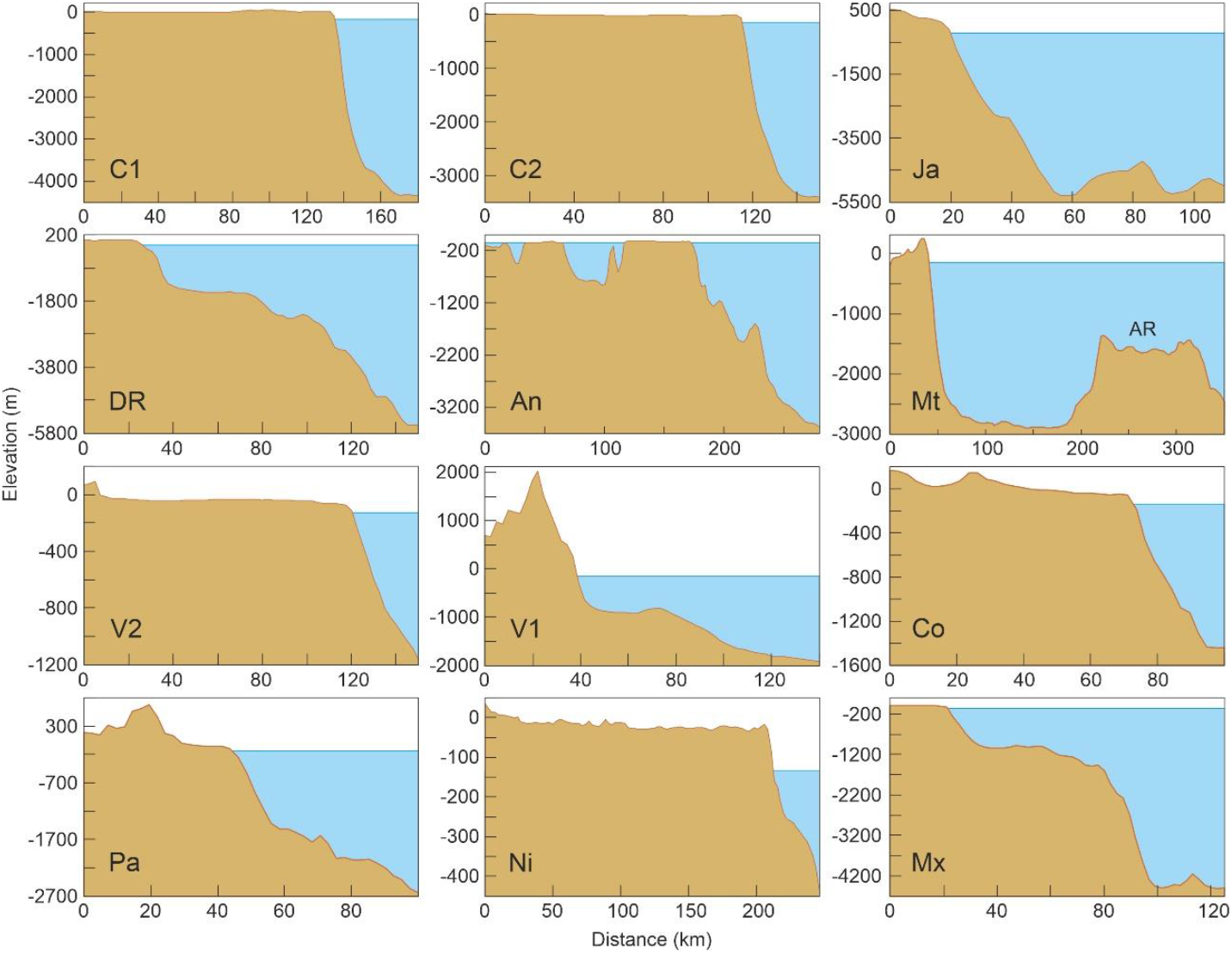
Distance-elevation profiles across the transects depicted in Fig. 3, showing the continental shelf/slope topography (brown areas) and the LGM sea level (blue) areas. Abbreviations: An, Anguilla; Co, Colombia; C1 and C2, Cuba; DR, Dominican Republic; Ja, Jamaica; Mt, Martinique (AR, Aves Ridge); Mx, Mexico; Ni, Nigaragua; Pa, Panama; V1 to V3, Venezuela.

### 3.2. Meso- and microtopography

The FA prospective sites identified in the mesotopographic analysis (Supplementary Material) were grouped into four main zones: the northern Trinidad shelf (NT), the Cariaco Basin (CB), the northern Colombian shelf (NC), and western Hispaniola (WH) (Fig. 3). The largest of these zones is NT, where the prospective FAs extend over approximately 200 km. The other zones consist of a few smaller spots, each less than 10 km wide. Along the remaining Caribbean coasts, the -128 to -133 m isobaths appear as thin lines, indicating steep coasts or coastal cliffs.

In the NT zone, FAs suitable for mangrove growth are more extensive in the central area, decreasing toward the east and west, where the coasts become steeper. This is illustrated by a central land-sea elevation profile (NTC), where the UML and LML span nearly 6 km inland with an average slope of 0.09% (Fig. 5). In contrast, an example of an unfavorable topographical gradient is found on the western side (NTW), where the UML and LML are less than 200 m apart, with an average slope of 3.05%. This slope is approximately 25 times the global average (0.12%) and seven times the maximum (0.45%) slope recorded worldwide for mangrove growth. A range of intermediate situations exists across the NT site, including slopes close to the maximum, as shown in an eastern profile (NTE) with an average slope of 0.41% between the UML and LML.

**Figure 5.**
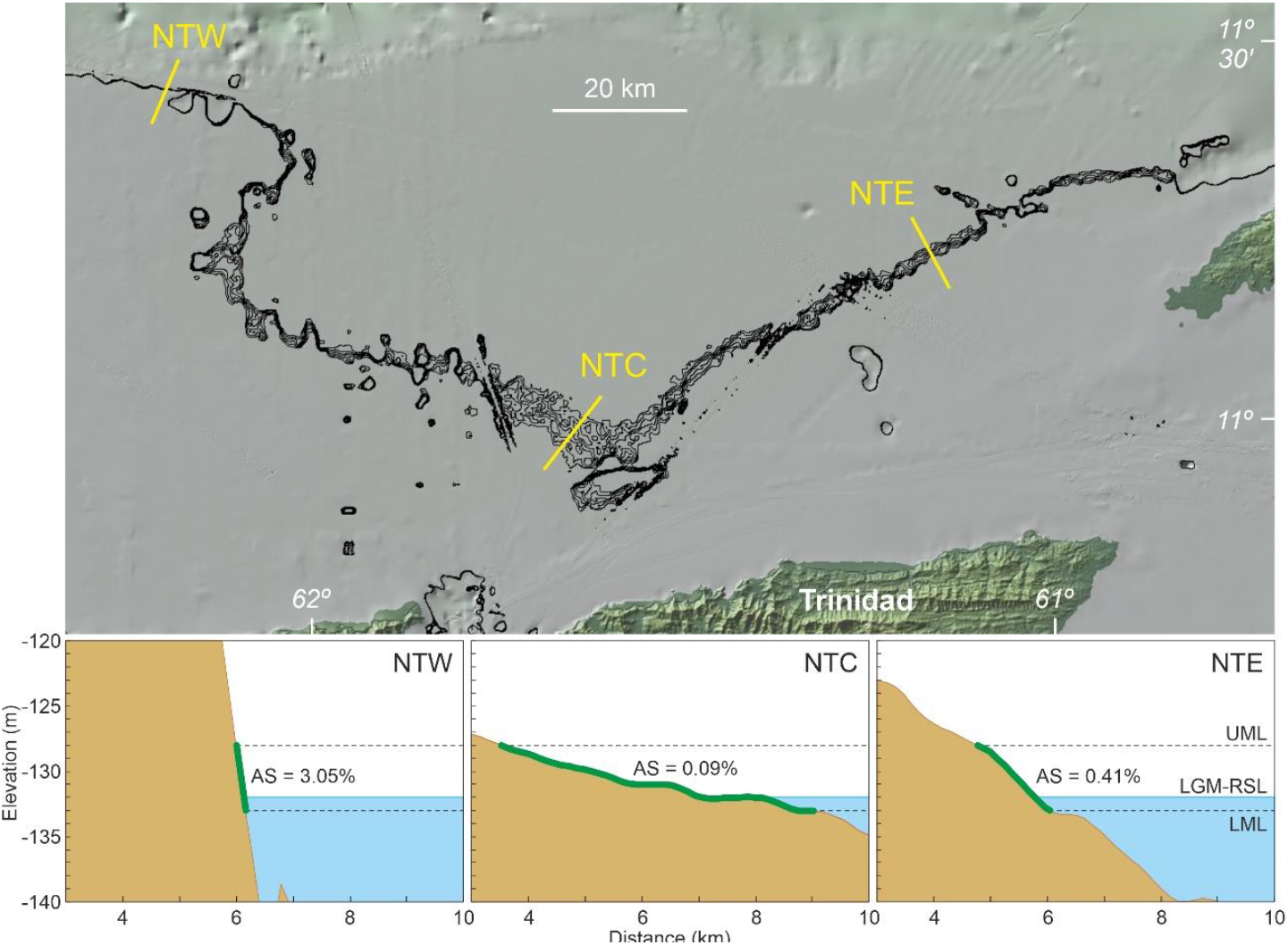
Close up of the NT site (Fig. 3) showing the isobaths between -128 and -133 m at meter resolution, and representative examples of LGM coastal profiles (yellow lines) from west to east. The space available for mangrove growth is highlighted by a thick green line in the below profiles. Abbreviations: AS, average slope; LGM-RSL, relative sea level during the Last Glacial Maximum; LML, lower mangrove limit (−133 m); UML, upper mangrove limit (−128 m).

In the CB zone, three small spots were identified, each within the slope range suitable for mangrove growth: 0.15% (CBN), 0.20% (CBS), and 0.24% (CBW) (Fig. 6). The remaining coasts in this zone appear as thin lines, resembling the steeper NTW profile in northern Trinidad. At the CBW site, the coastal profile is more complex than a single slope, and certain areas, particularly in the center of the profile, may be too steep for mangrove growth.

**Figure 6.**
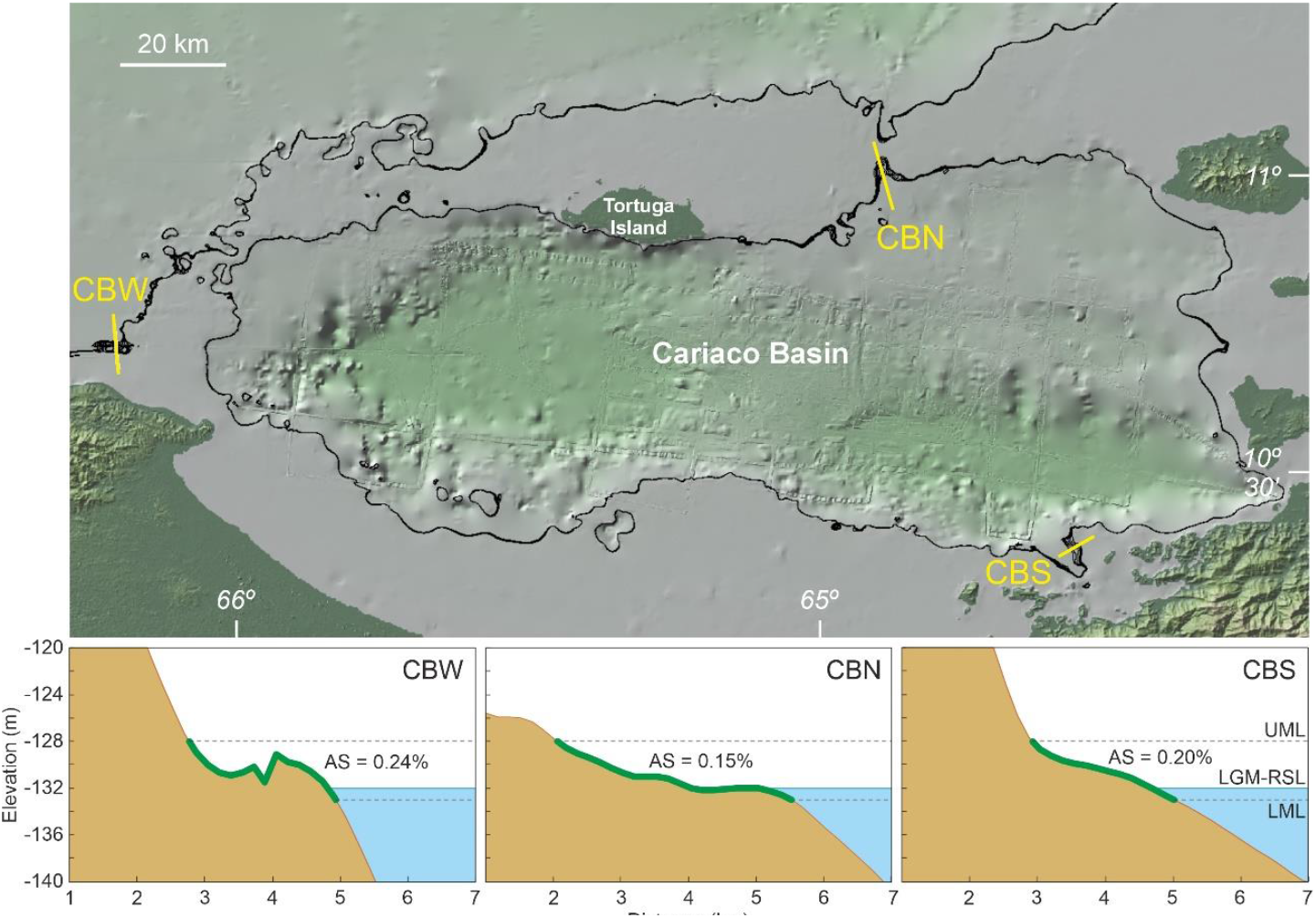
Close up of the CB site (Fig. 3) showing the isobaths between -128 and -133 m at meter resolution and the land-sea profiles cross the prospective sites identified (yellow lines). The space available for mangrove growth is highlighted by a thick green line in the below profiles. Abbreviations: AS, average slope; LGM-RSL, relative sea level during the Last Glacial Maximum; LML, lower mangrove limit (−133 m); UML, upper mangrove limit (−128 m).

In the NC zone, two prospective sites were identified. The flatter site is located in the south (NCS; 0.17%), while the steeper site, though still within the mangrove range, is in the north (NCN; 0.28%) (Fig. 7). The WH zone presents a unique case. While the profile meets the conditions for mangrove growth, with slopes ranging from 0.23% in the northwest to 0.36% in the southeast, it is a closed basin—likely a coastal lagoon—of unknown water salinity.

**Figure 7.**
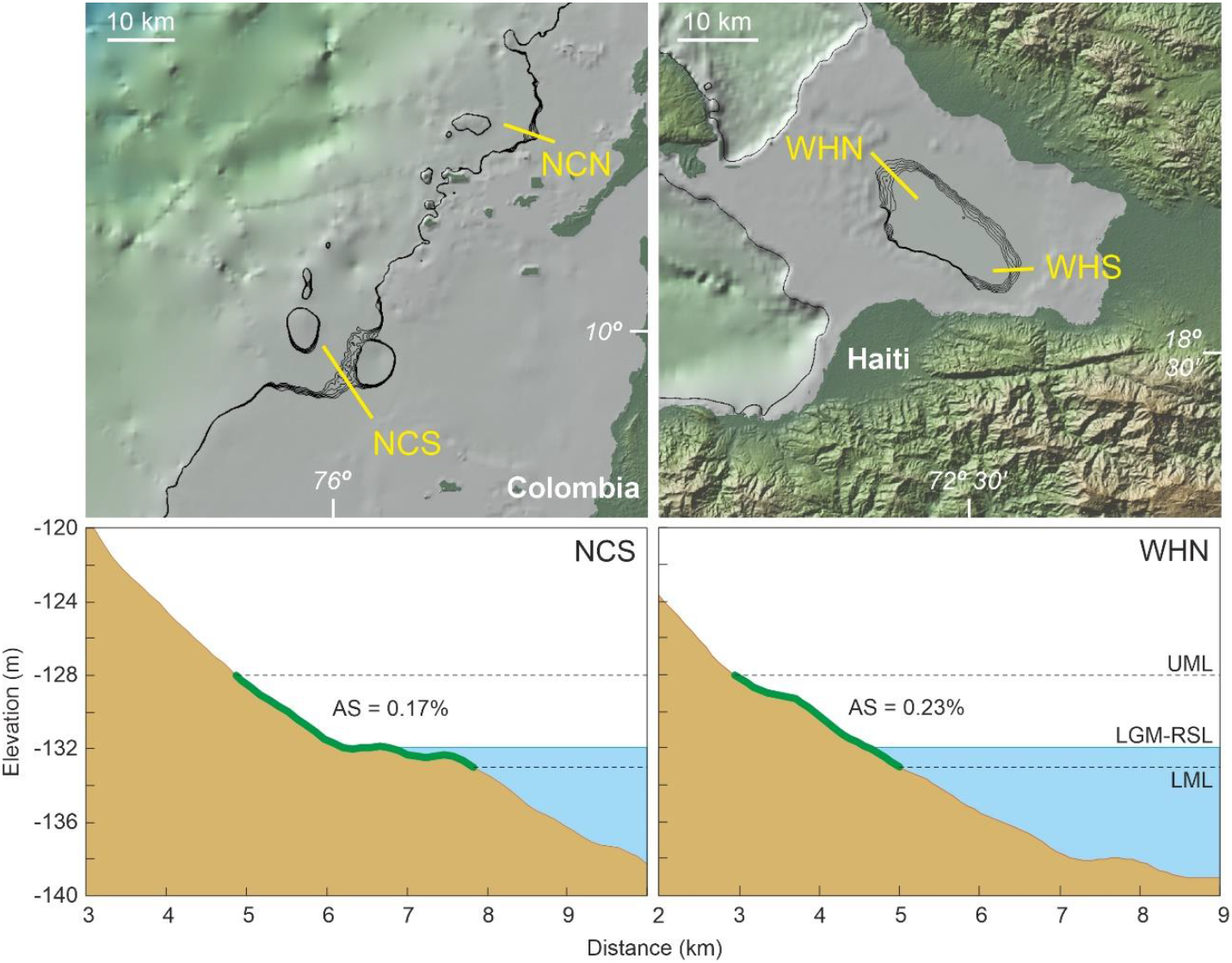
Close up of the NC and WH zones (Fig. 3) showing the isobaths between -128 and -133 m at meter resolution and the land-sea profiles across the prospective sites identified (yellow lines). The space available for mangrove growth is highlighted by a thick green line in the below profiles. Abbreviations: AS, average slope; LGM-RSL, relative sea level during the Last Glacial Maximum; LML, lower mangrove limit (−133 m); UML, upper mangrove limit (−128 m).

## Conclusions and discussion

The results indicate that during the LGM, most Caribbean coasts were unsuitable for mangrove growth due to the aerial exposure of flat continental shelves and the steep land-sea elevation gradients typical of continental slopes. However, some zones remained where mangroves could have survived, considering the characteristic topography associated with these ecosystems. The largest accumulation of mangroves likely existed on the northern Trinidad shelf (NT), which may be considered the primary LGM mangrove refugium. Other topographically suitable areas consisted of small (km-scale) and scattered sites that fall under the category of microrefugia (sensu Rull, 2009).

Bioclimatically, these potential refugia were suitable for mangrove growth, despite the LGM cooling and hydroclimatic reorganizations. At present, the average air temperatures along the coasts adjacent to these refugia range from approximately 25°C to 28°C (Table 1). Given an estimate Neotropical (30°N to 30°S) cooling of 5-6°C below present temperatures (Bush et al., 2001) and adjusting by the 132-m RSL drop using a conservative temperature lapse rate range of -0-5°C to -0.8°C per 100 m elevation (Martin & Fahey, 2014), the average LGM temperatures in the potential mangrove refugia would have ranged from approximately 20°C to 24°C. These temperatures remain well above the lower average limits of the major components of Caribbean mangroves, such as *Rhizophora* (19.2°C) and *Avicennia* (13.5°C) (Quisthoudt et al., 2012). Therefore, temperature was not a limiting environmental driver for mangrove growth.

**Table 1.** Climatic data (1991-2020) from coastal areas adjacent to the identified LGM refugia for Caribbean mangroves. MAP, mean total annual precipitation; MAT, mean annual temperature. Raw data from the Climatic Research Unit (CRU) of the University of East Anglia (https://www.uea.ac.uk/groups-and-centres/climatic-research-unit), mapped by the World Bank Group (https://climateknowledgeportal.worldbank.org/; last accessed March 11, 2025).

|  | NT | CB | NC | WH |
| --- | --- | --- | --- | --- |
| **MAT (°C)** | 26.3-26.6 | 25.4-27.6 | 28.2 | 24.5 |
| **MAP (mm)** | 1398-2043 | 746-1056 | 1677 | 1466 |

Regarding total annual precipitation, which currently ranges between approximately 750 mm and 2050 mm along the neighboring coasts (Table 1), Caribbean-wide LGM estimates are hindered by the high spatial variability observed in paleoecological records (Warken et al., 2019). However, precipitation was also unlikely to be a limiting factor, as mangroves thrive across a wide range of hydroclimatic conditions worldwide, from arid (<100 mm/yr) to very humid (>3000 mm/yr) environments (Osland et al., 2017). The development of environmental niche modeling (ENM) studies (Thuiller, 2024) is beyond the scope of this paper and is unlikely to alter the regional bioclimatic framework for mangrove occurrence along the Caribbean coasts during the LGM. However, these studies could provide more detailed insights into each potential refugium identified in this paper.

Interestingly, the areas suitable for mangrove growth during the LGM were concentrated along the South American Caribbean coasts, particularly in the southeastern Caribbean region (northern Trinidad and northeastern Venezuela). No suitable FAs were identified along the remaining coasts, except for the western Hispaniola site, which was likely a coastal lagoon. This suggests that during the postglacial sea-level rise, mangroves may have colonized the entire Caribbean coastline primarily from refugia and microrefugia located in the southeastern region. Given that surface coastal currents are the main agents for dispersing mangrove propagules, especially in the Caribbean and other tropical regions worldwide (Van der Stocken et al., 2019), understanding the surface circulation patterns during the LGM is essential to evaluate the consistency this hypothesis, though not the hypothesis itself.

Currently, surface currents enter the Caribbean through the eastern coasts via the Northern Equatorial Current (NEC) and the Guyana Current (GC). The first flows through the Lesser Antilles, while the second enters through northern Trinidad and the Cariaco Basin (Fig. 8). Inside the Caribbean Sea, these currents form the Caribbean Current (CC), which flows in an ESE-NW direction before entering the Gulf of Mexico through the Yucatan Channel (Giry et al., 2013; Osborne et al., 2014; Felis et al., 2015). Isotopic evidence and modeling suggest that during the postglacial and Holocene sea-level rise, the general circulation patterns remained similar to those observed today, with minor variations in intensity and seasonality (Giry et al., 2013; Gu et al., 2017). Therefore, surface circulation would have been favorable for westward mangrove expansion from the primary LGM refugial areas in northern Trinidad and the Cariaco Basin across the Caribbean. Minor contributions from other microrefugia are also consistent with this circulation pattern.

**Figure 8.**
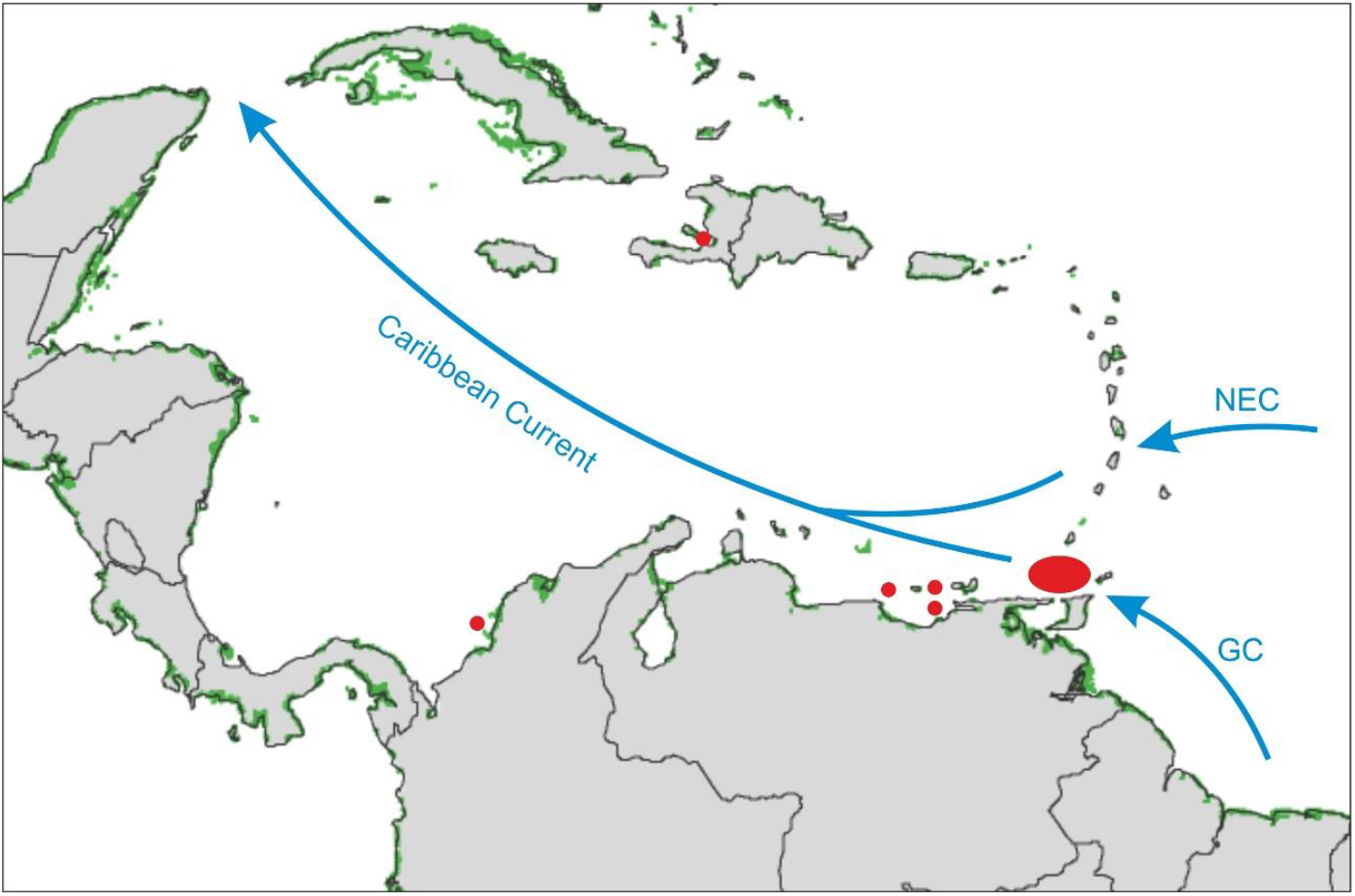
Map showing current mangrove areas as in Fig. 1 (green patches), the LGM refugial areas identified in this work (red dots) and the main superficial Caribbean currents (blue arrows). See text for more details. NEC, Northern Equatorial Current; GC, Guyana Current.

It should be emphasized that the occurrence of FAs around -132 m, as identified in this study, represents a minimum requirement for LGM mangrove persistence. Other critical environmental factors for mangrove growth, such as salinity, river discharge, and sediment/nutrient supply, can only be determined through paleoecological surveys involving marine coring. This study focuses on identifying the most promising paleoecological prospects, significantly reducing the area for potential LGM mangrove investigations and saving considerable time, effort, and resources.

According to the study results, the highest priority should be given to the northern Trinidad shelf, especially its central area (Fig. 5), which hosts the most extensive refugial site identified. This region could have been the primary inoculation zone for modern Caribbean mangroves following the LGM bottleneck that severely reduced mangrove extent. Coring in the NT prospect and nearby offshore sediments is strongly recommended to elucidate mangrove dynamics during the LGM. The final selection of specific coring targets should be guided by seismic surveys of Pleistocene sediments. The proximity of gas fields and the associated facilities and infrastructures (NGC, 2023) could be of help in this sense.

The Cariaco Basin is a classical Neotropical site for Quaternary paleoclimate and paleoenvironmental reconstruction and has been intensively studied and cored. However, LGM sediments, if present, have not yet been analyzed—or results have not been published—for vegetation reconstruction and mangrove occurrence. The scarcity of mangrove pollen in slightly older sediments (68–28 ka BP; González et al., 2008; González & Dupont, 2009) may reflect the microrefugial nature of the surrounding sites (Fig. 6). Nonetheless, palynological analysis of potential LGM intervals in these sediments is recommended. Other potential microrefugial areas, such as NC and WH, also merit coring. In the NC zone, deeper offshore sediments should be targeted, while in WH, the best prospect appears to be the lagoonal depocenter. As with NT, seismic profiles would aid in identifying precise coring targets.

With the current information, alternative mangrove sources for colonization outside the Caribbean cannot be ruled out. For instance, the Guyana Current may have transported propagules from the Atlantic coasts; however, the existence of LGM mangrove refugia along these coasts remains unconfirmed. The Pacific coasts of Central America are separated from the Caribbean by a land barrier that would have been difficult for littoral ecosystems to cross, particularly under lowstand conditions. The Gulf of Mexico is also an unlikely source for Caribbean mangroves, as surface currents flow in the opposite direction. These hypotheses require further study to establish a consistent regional framework. In addition to paleoecological research, phylogenetic studies of extant mangroves could help inferring the geographical origin of current Caribbean mangroves.

## Acknowledgments

No financial support was required specifically for the development of this work. The author is grateful to Andrew Goodwille for recommending the use of the GMRT tool for image analysis. The CERCA Programme, Generalitat de Catalunya, is also acknowledged.

## Conflict of interest

The authors declare no conflict of interest.

## Data availability

The data for this study are displayed in the manuscript.

## Notes

### Competing Interest Statement

The authors have declared no competing interest.

### Summary of Updates

The estimated bioclimatic conditions of the identified LGM refugia have been added, to support their suitability for mangrove growth.

