## Supporting material for "Where were the Caribbean mangroves during the Last Glacial Maximum? A microtopographical approach" (from another version of this preprint: version 2, 2025-01-31)

Valentí Rull

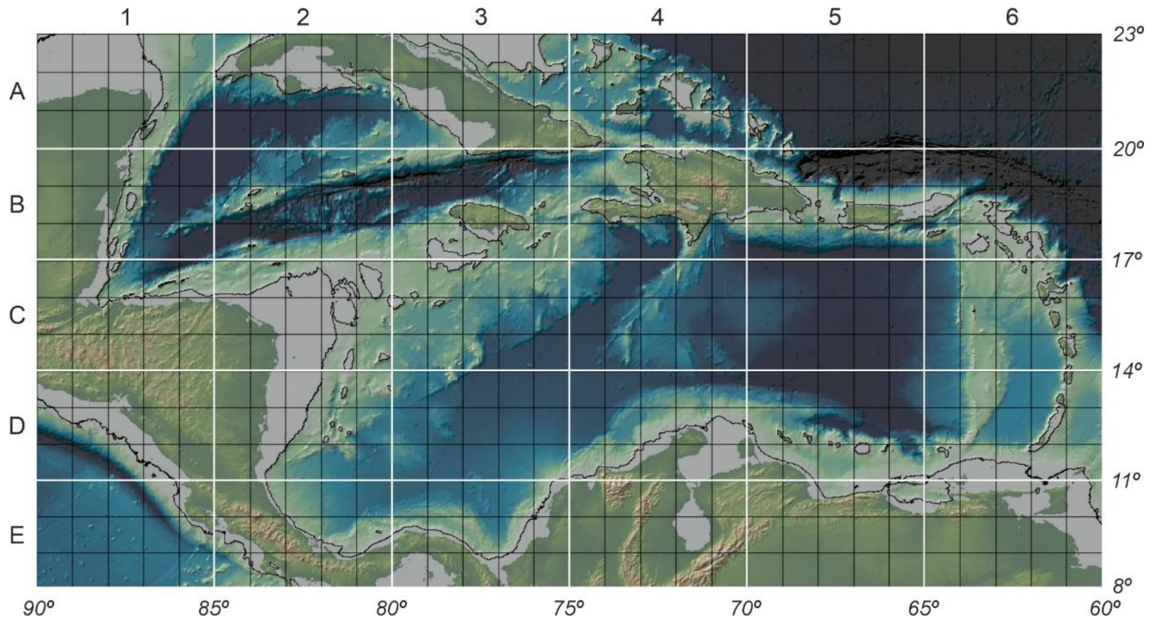


**Figure S1.** Reference map for the mesotopographical survey indicating the 30 arbitrary subdivisions selected. The -128 to -133 m isobaths are indicated. Only the blocks containing Caribbean coasts have been magnified in the high-resolution study (see below). Therefore, the following blocks have not been considered: A5, A6, C4, C5, D1, D3, E1 and E4. Prospective sites to be analyzed with more detail in the microtopographical survey are indicated with a yellow arrow. See the methods section for more details. Base map as in Fig. 2 of the main text.

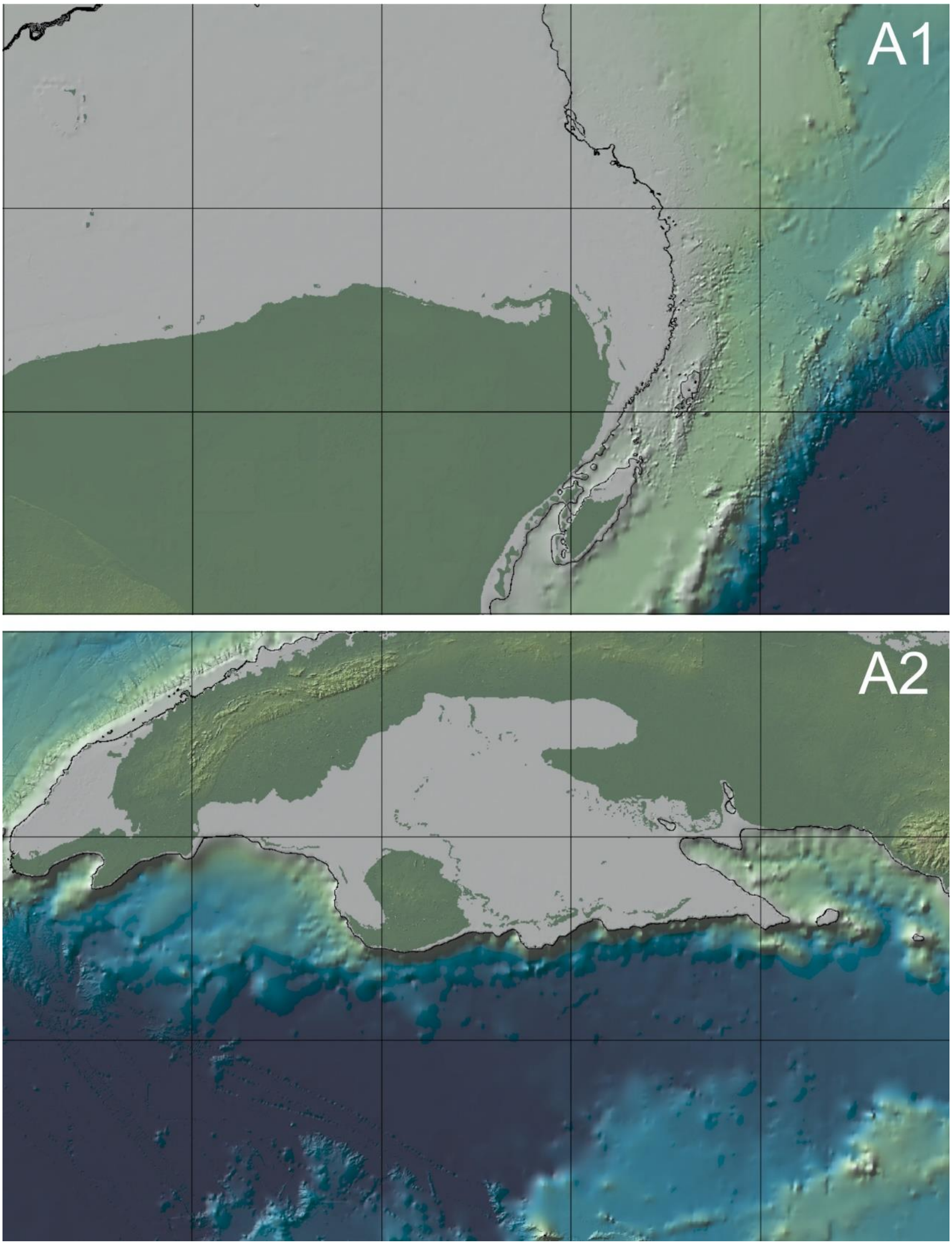
A1
A2

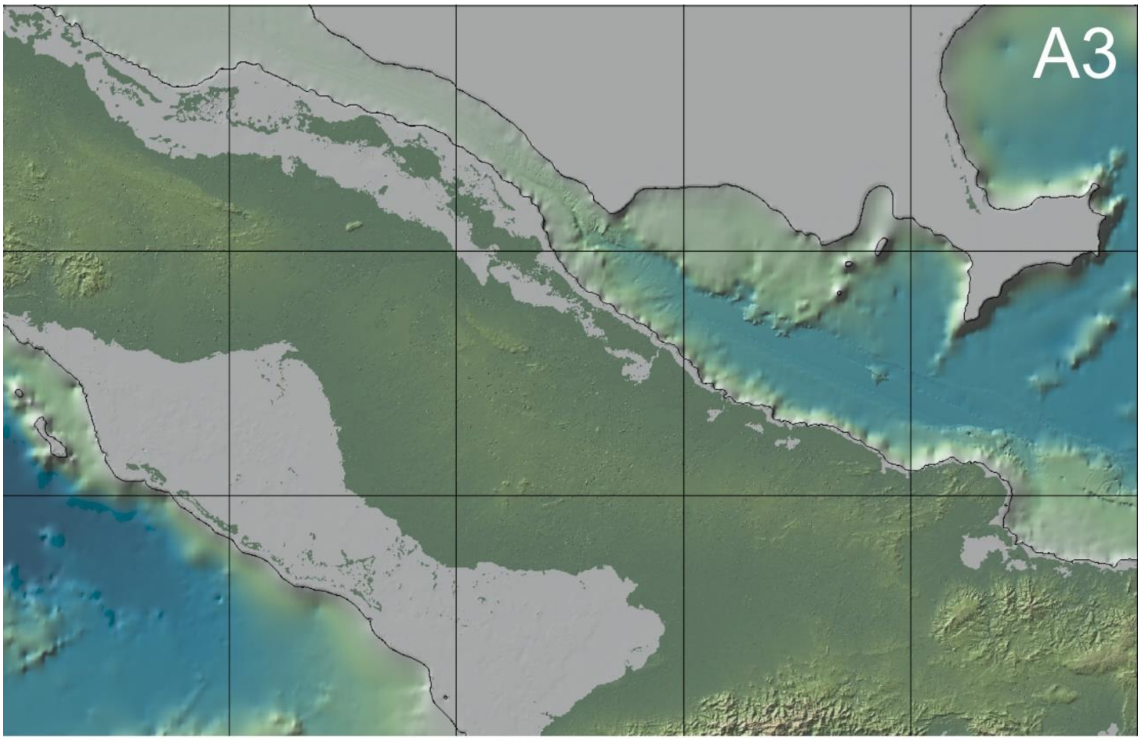
A3

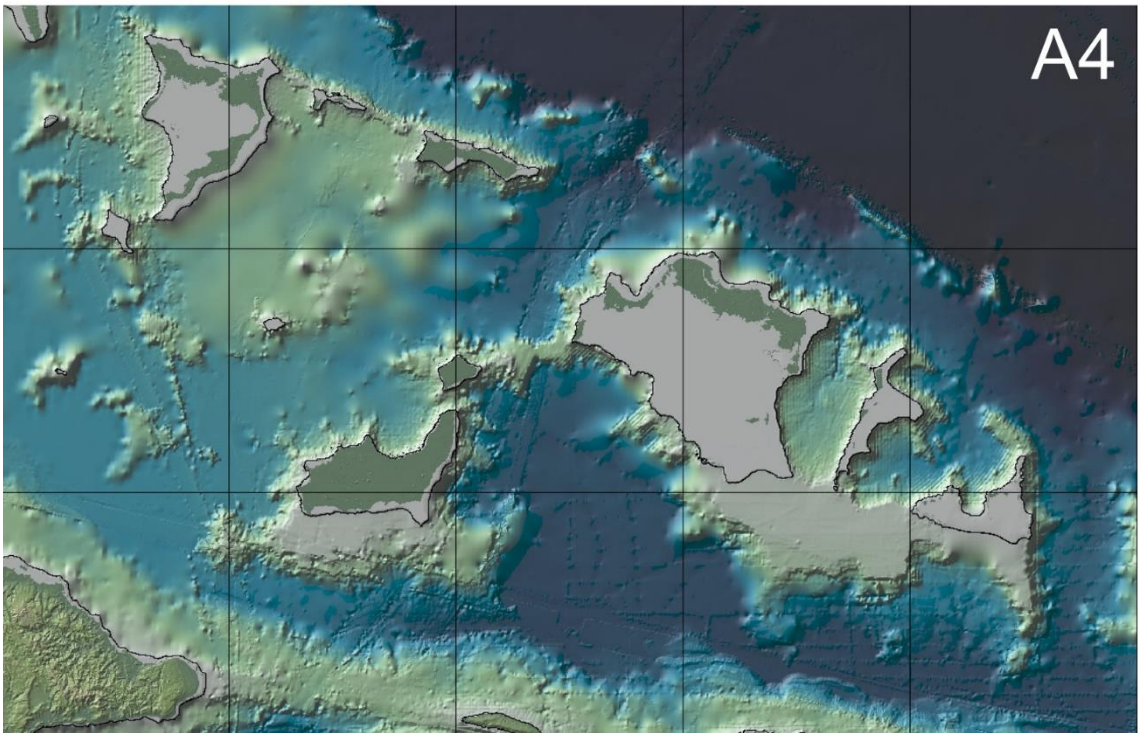
A4

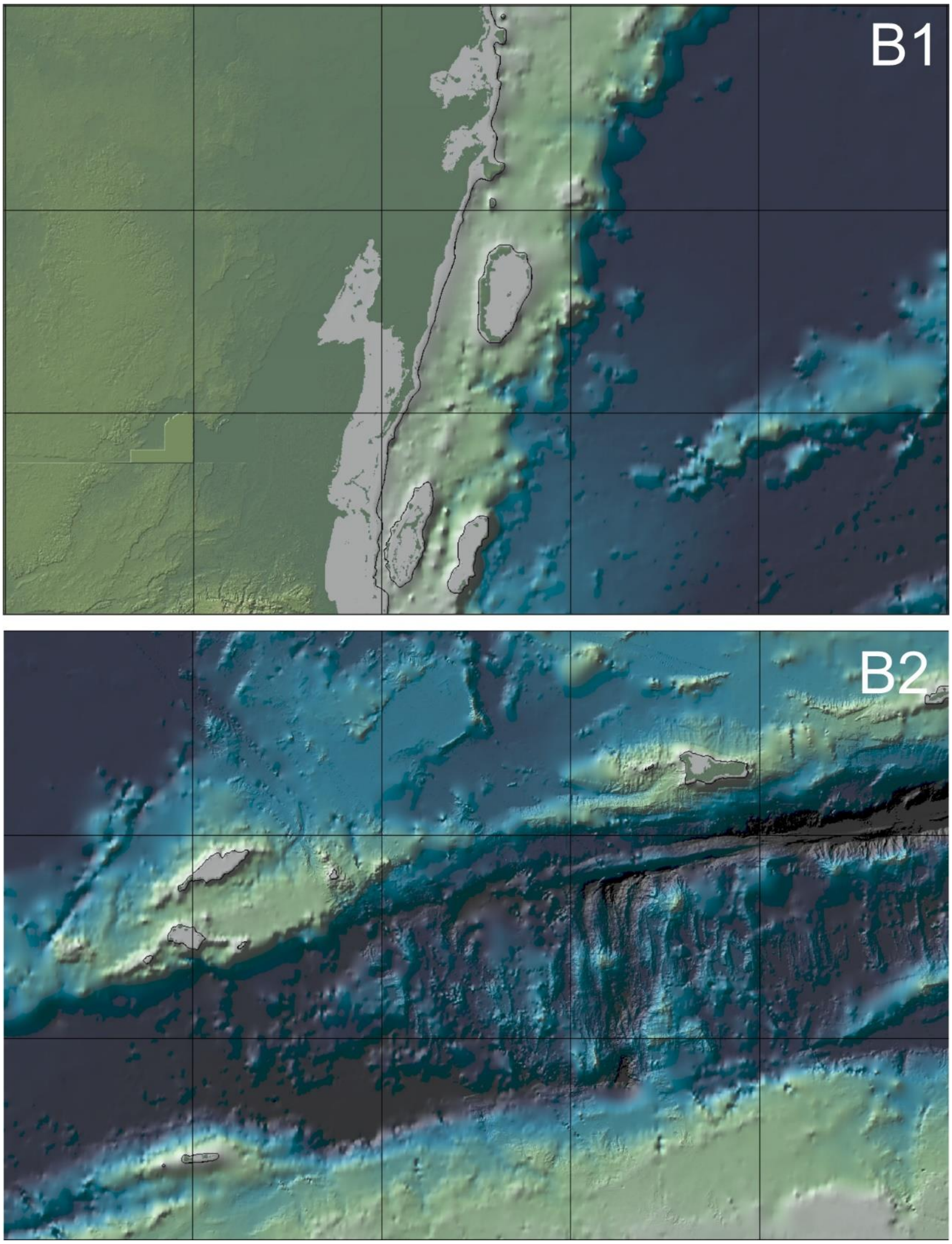
B1
B2

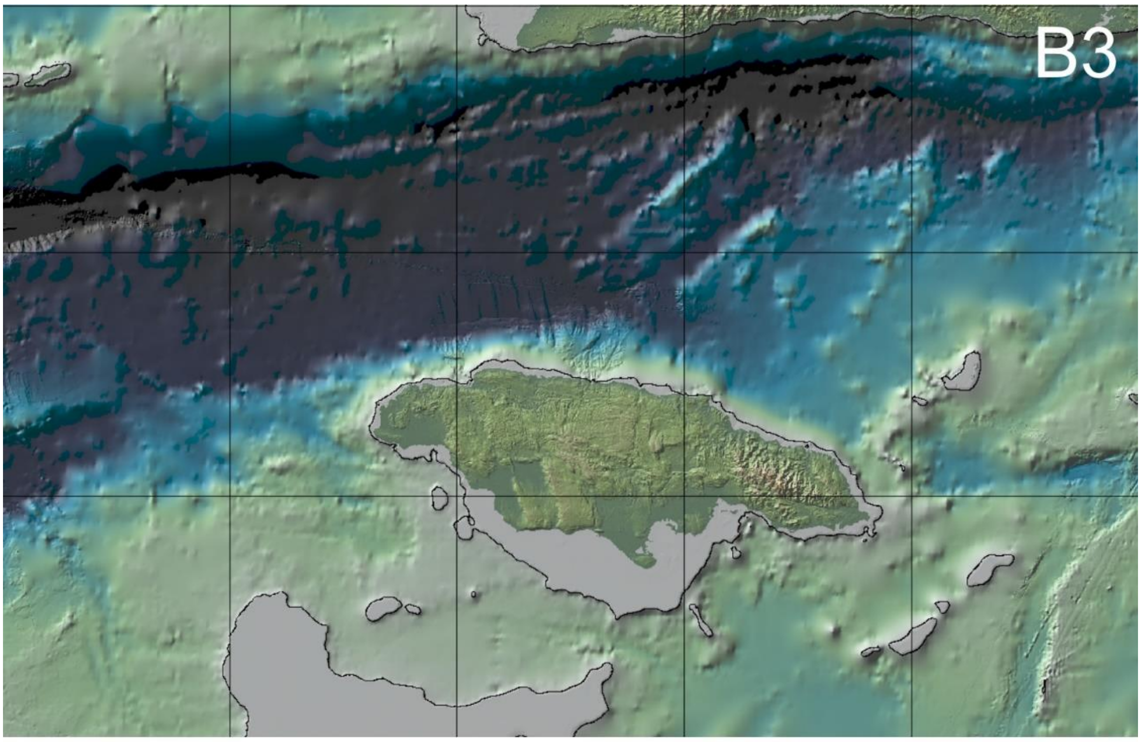
B3

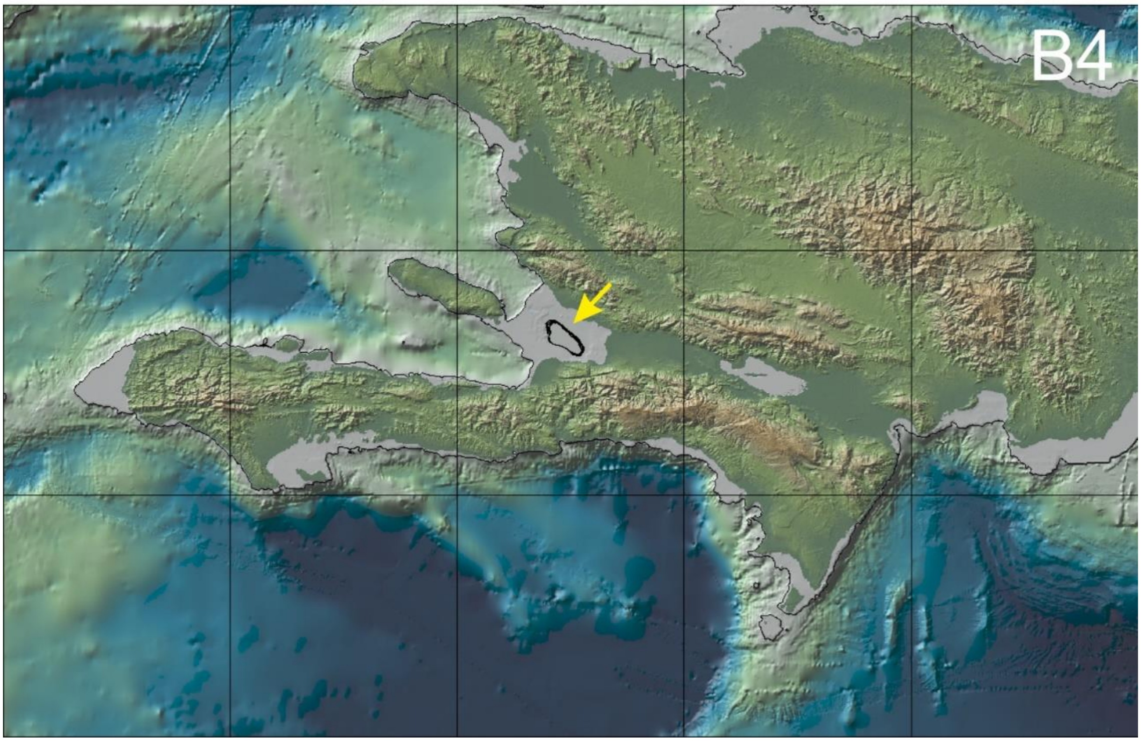
B4

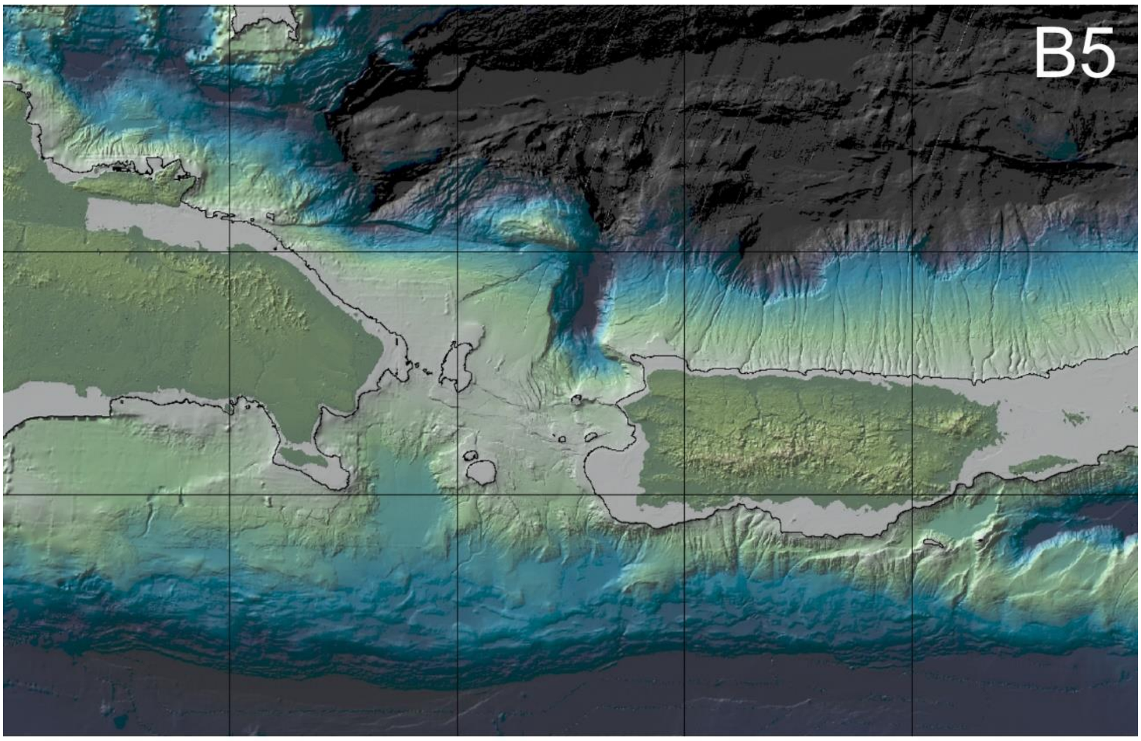
B5

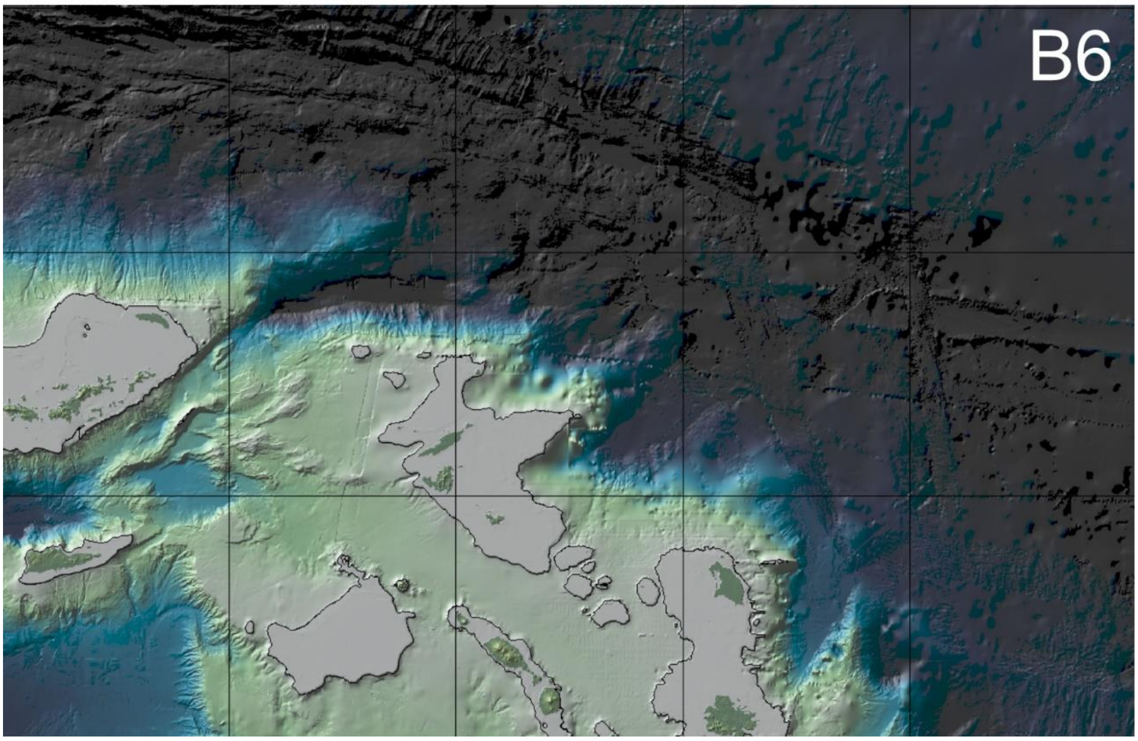
B6

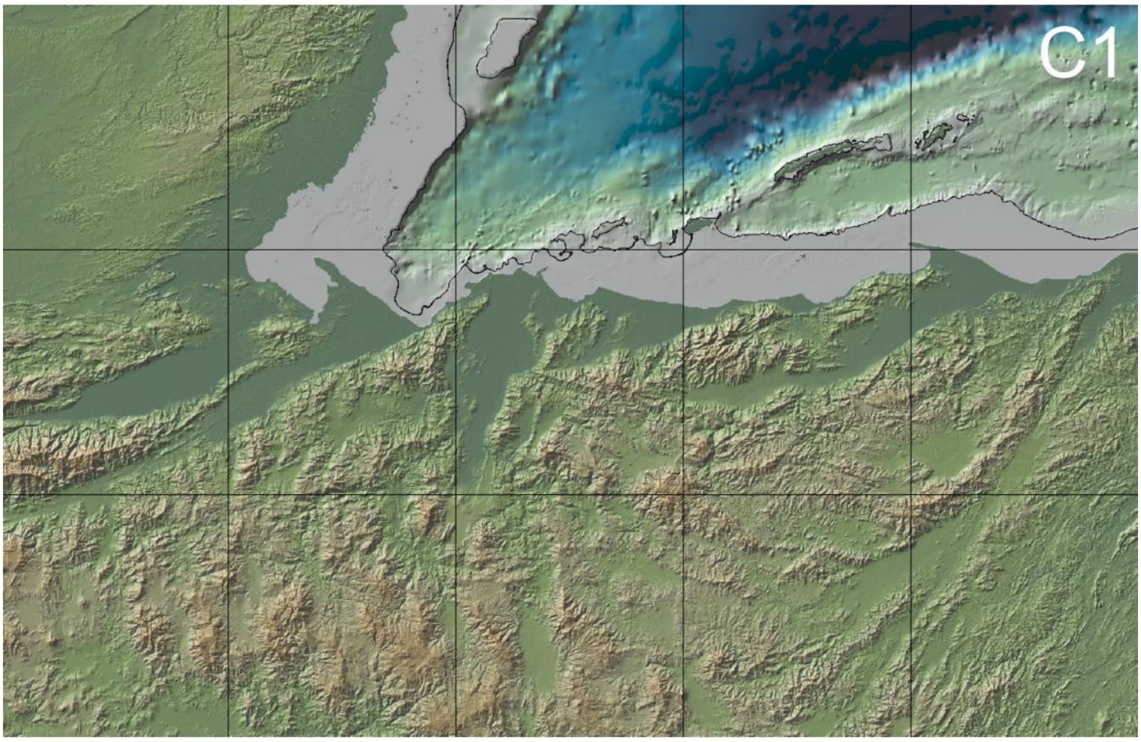
C1

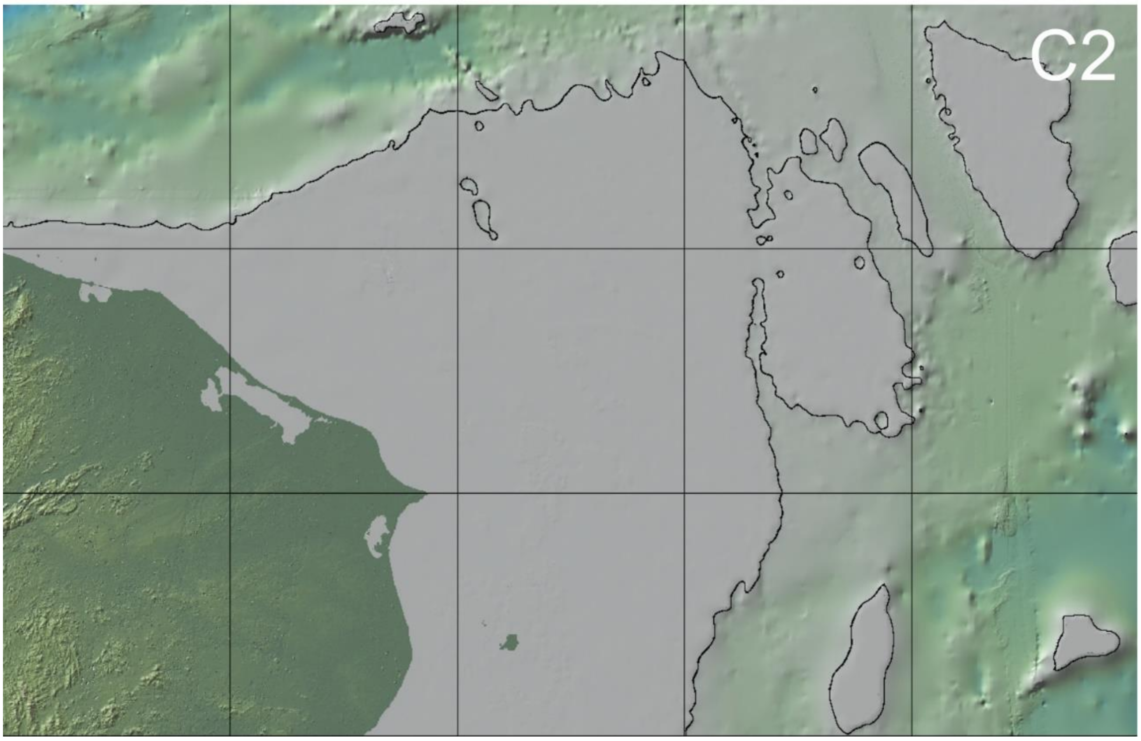
C2

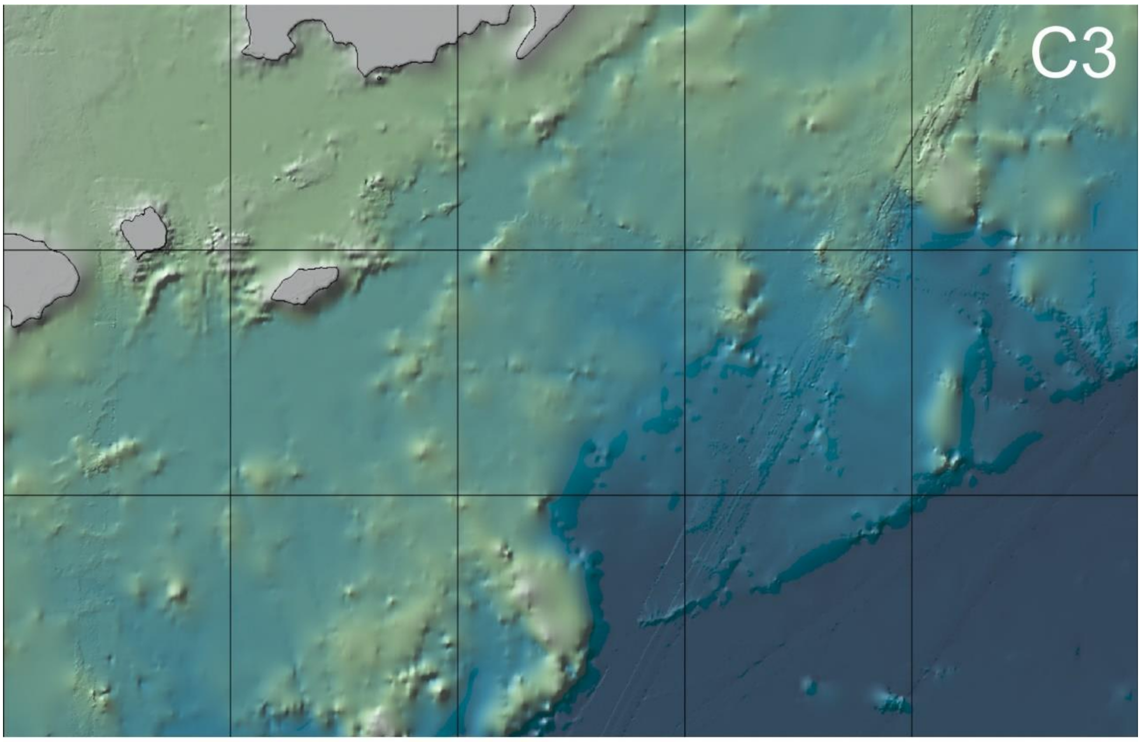
C3

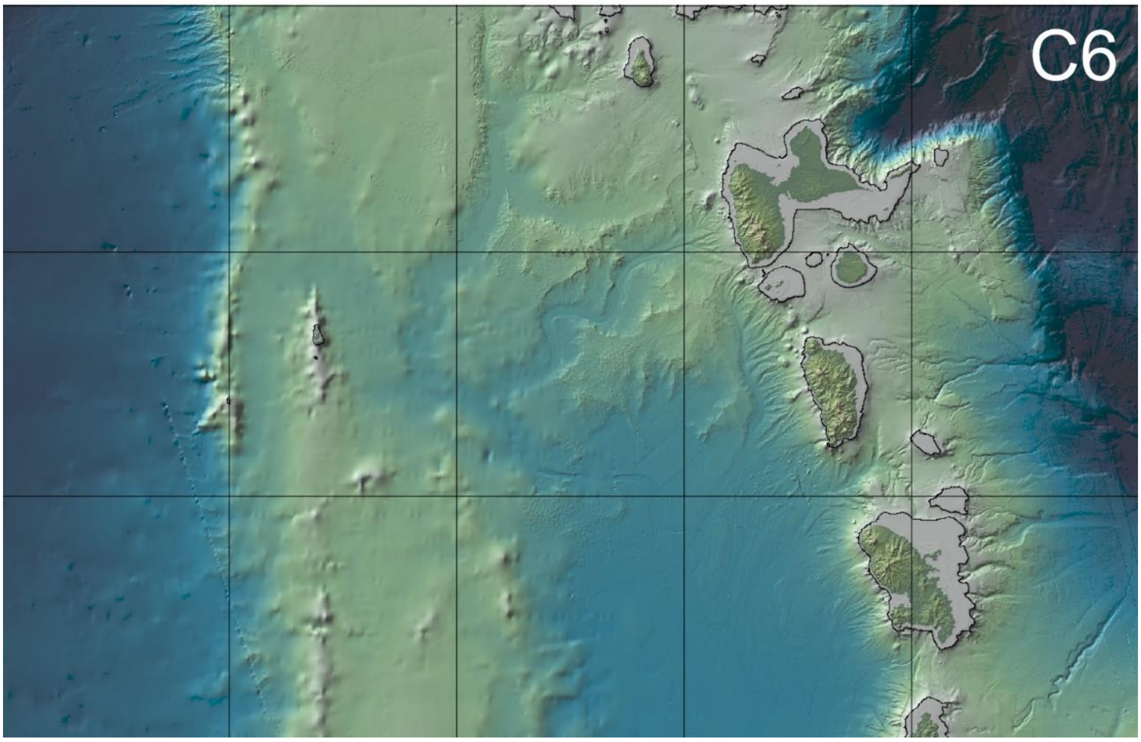
C6

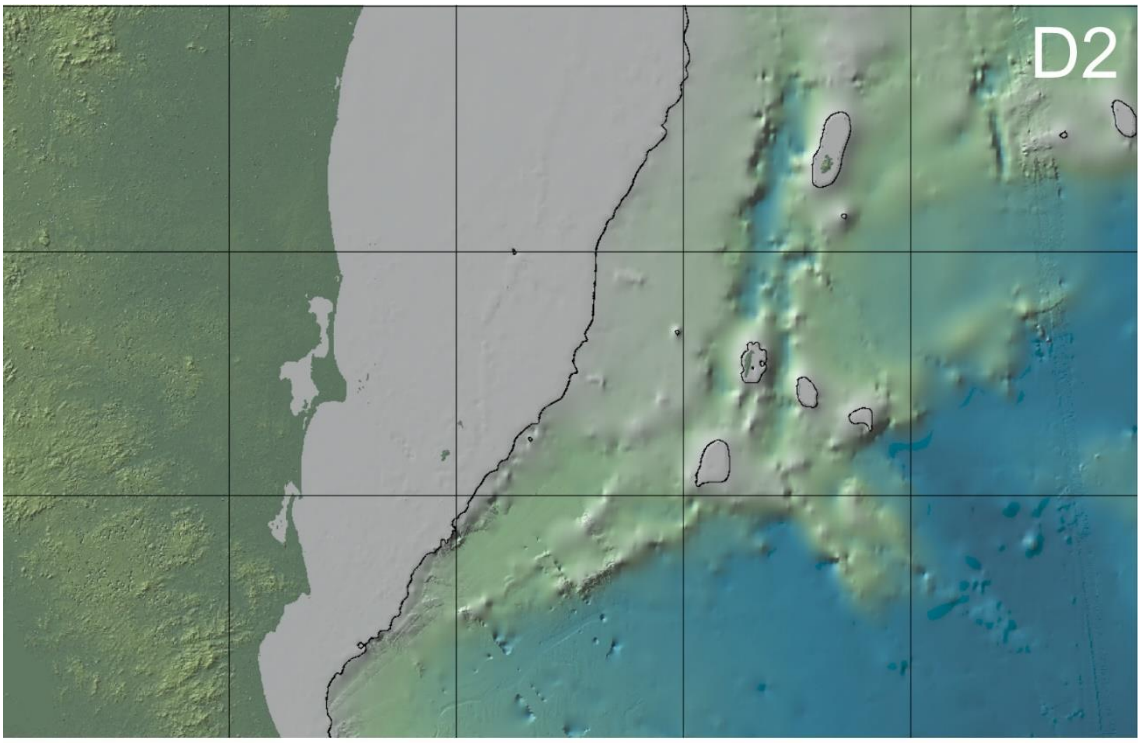
D2

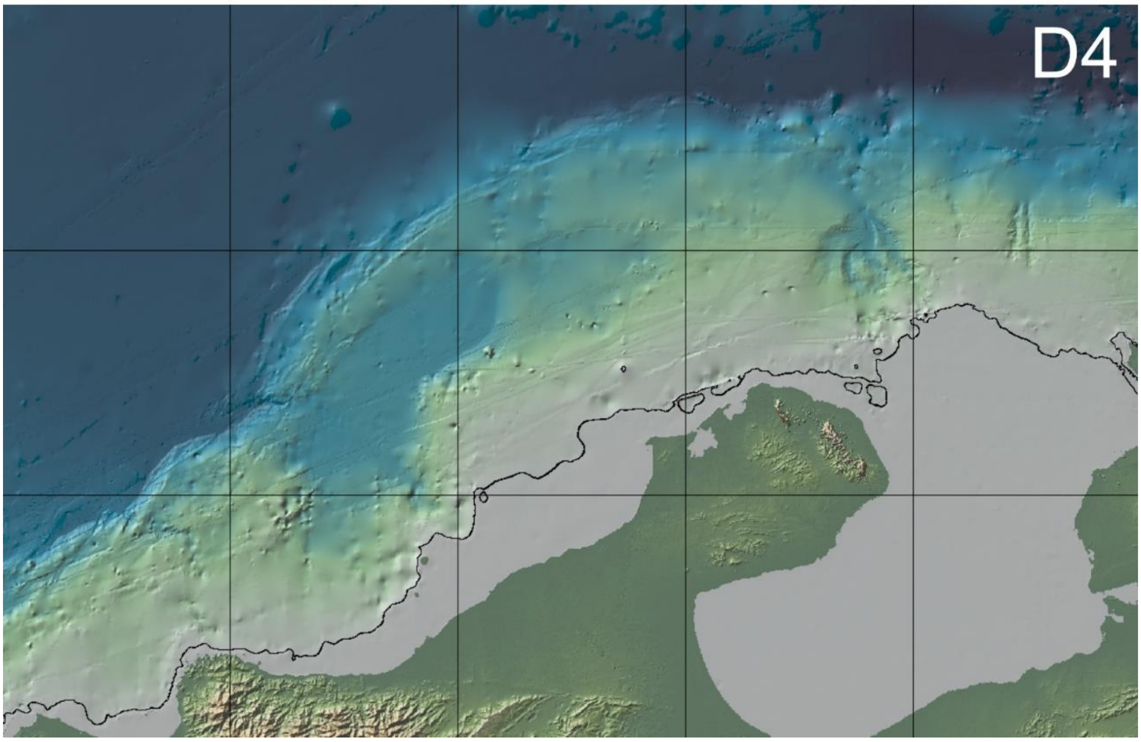
D4

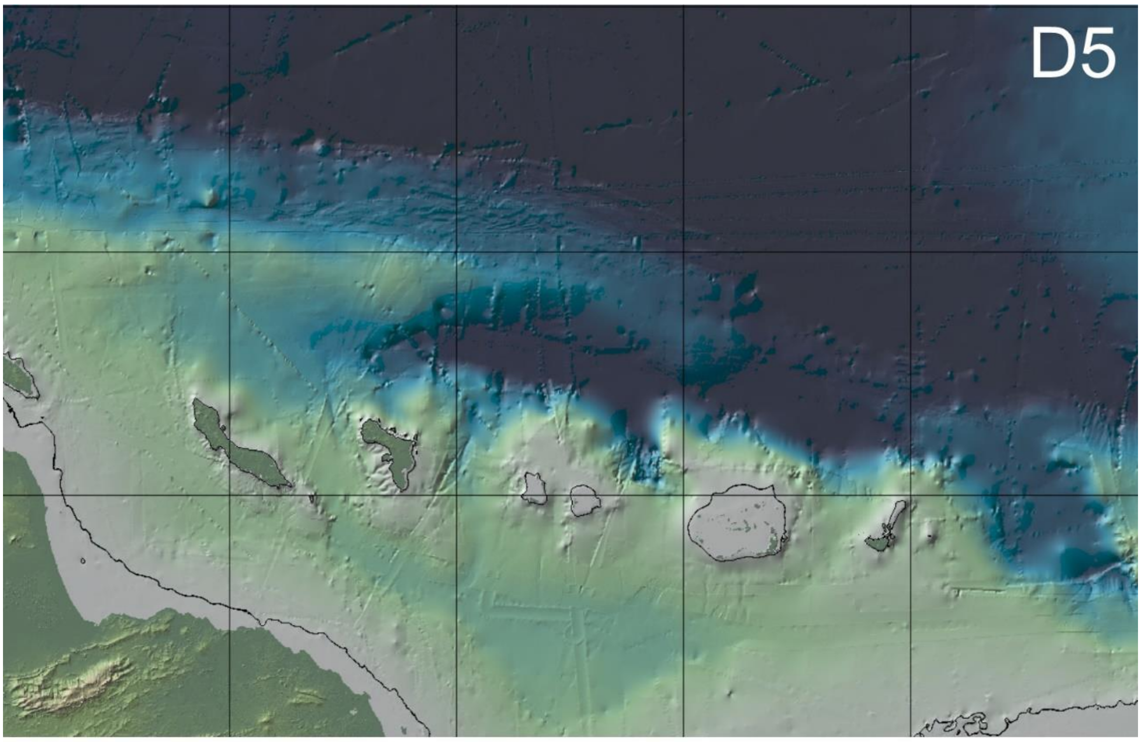
D5

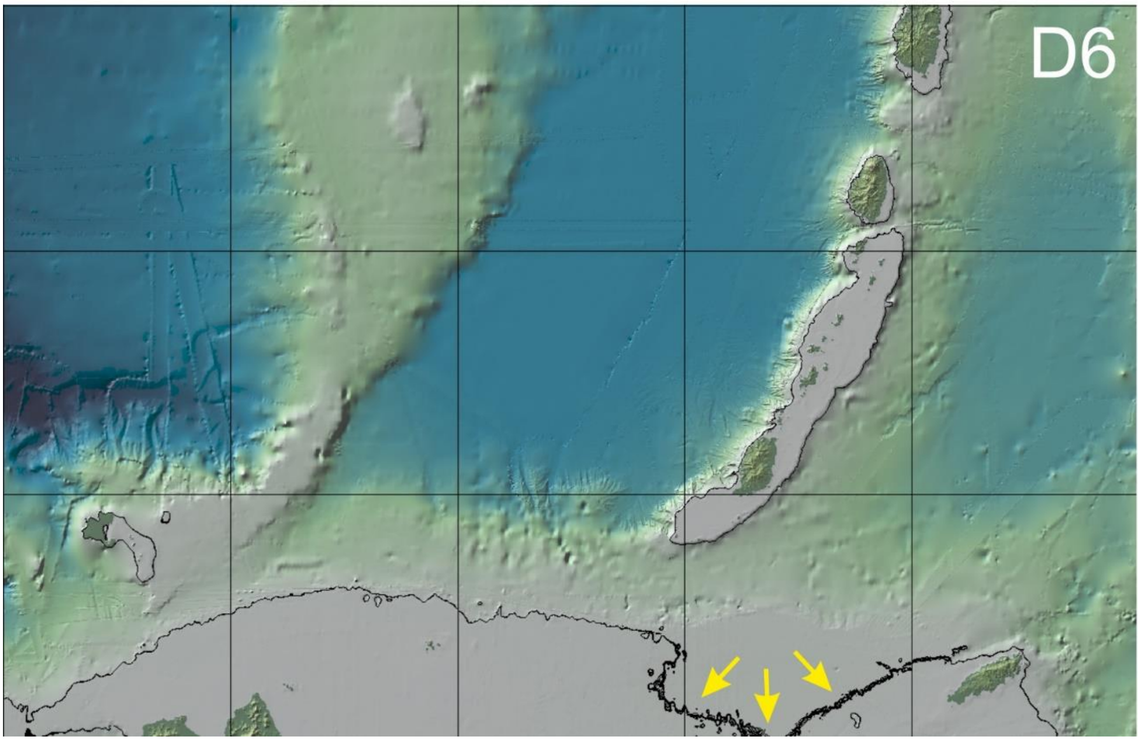
D6

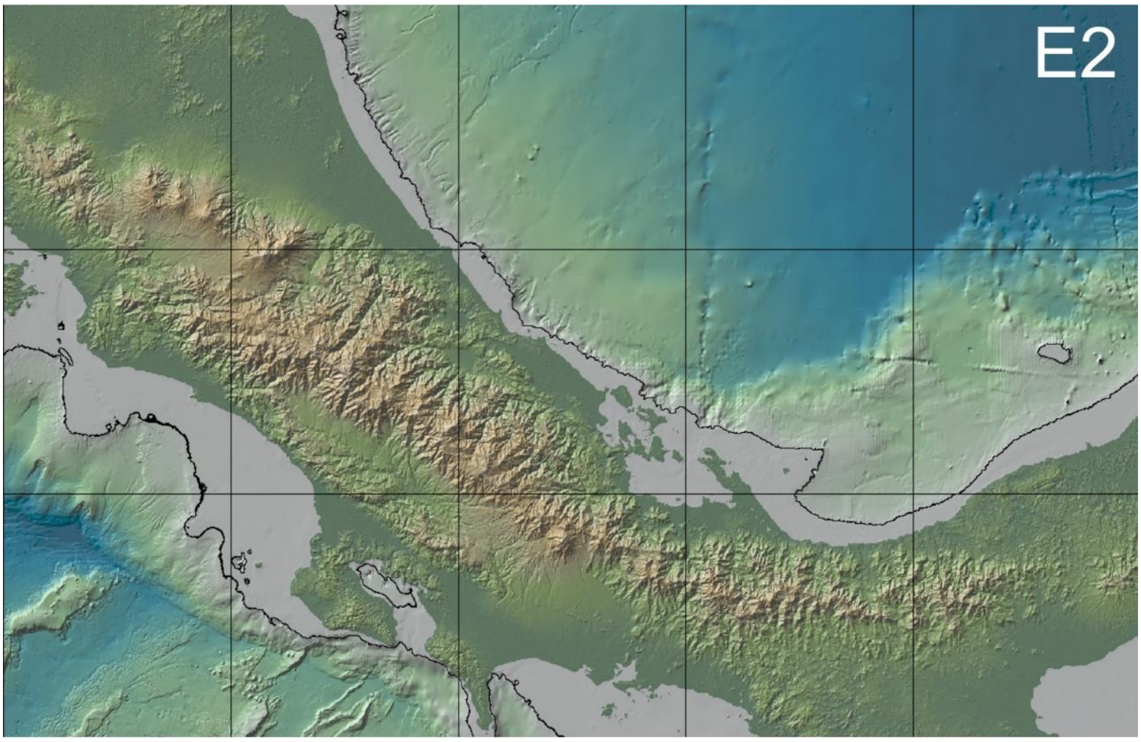
E2

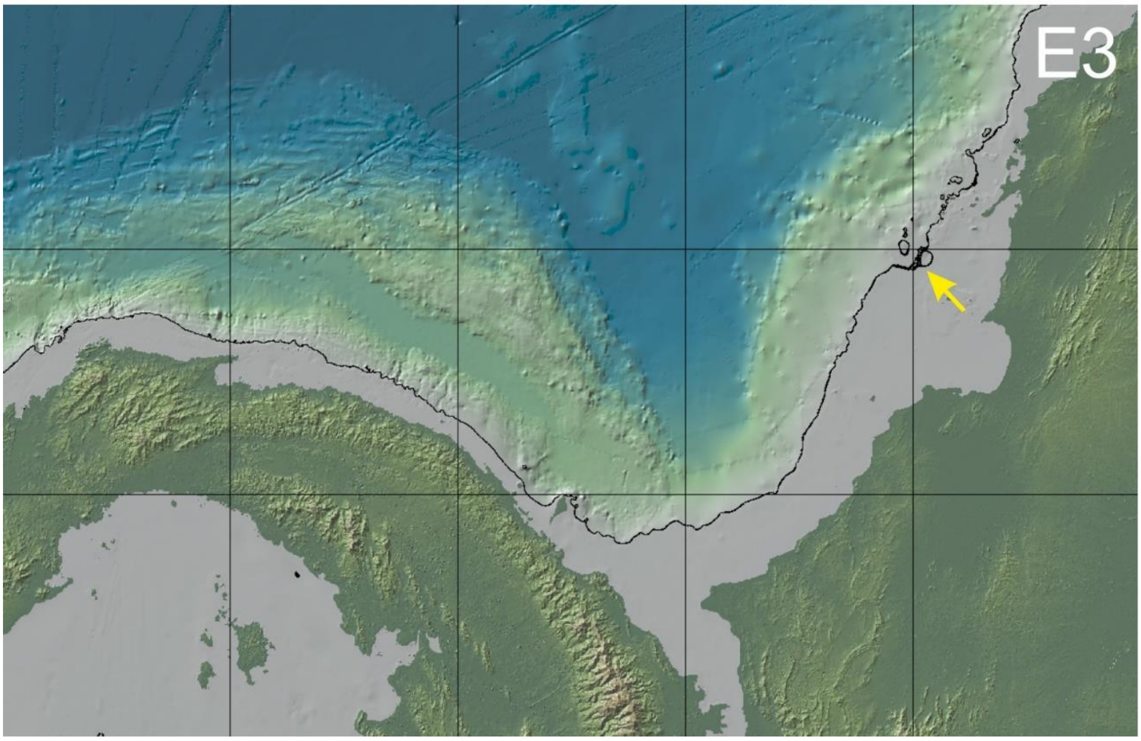
E3

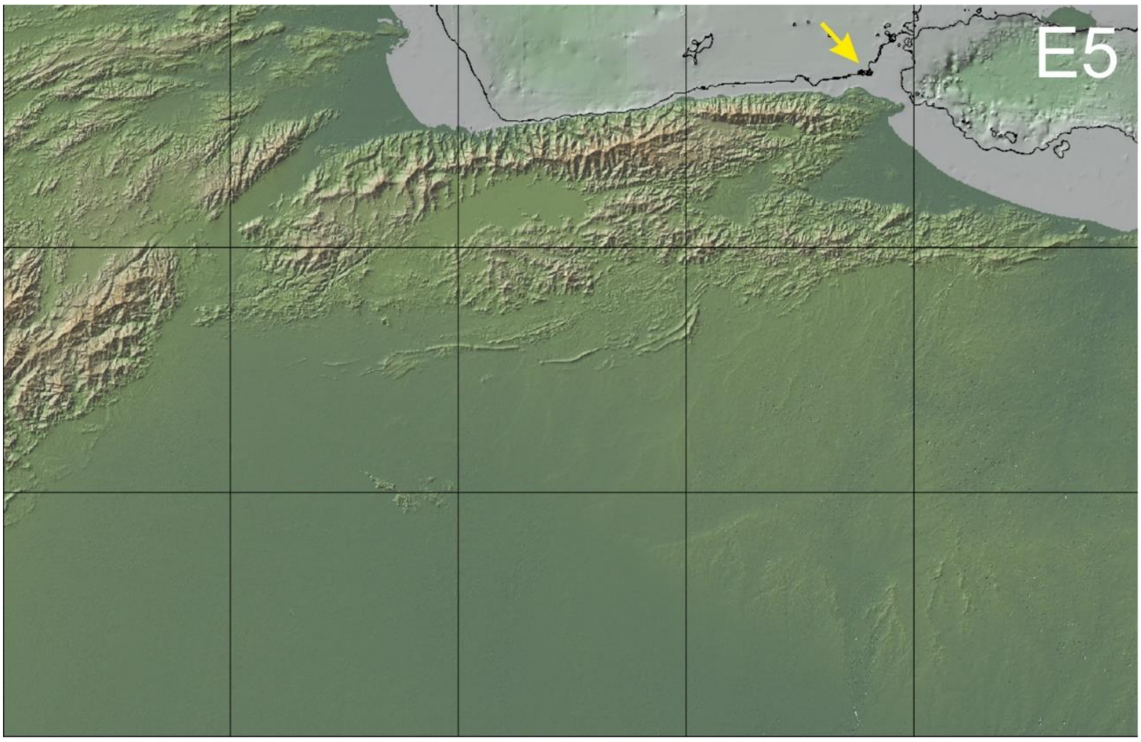
E5

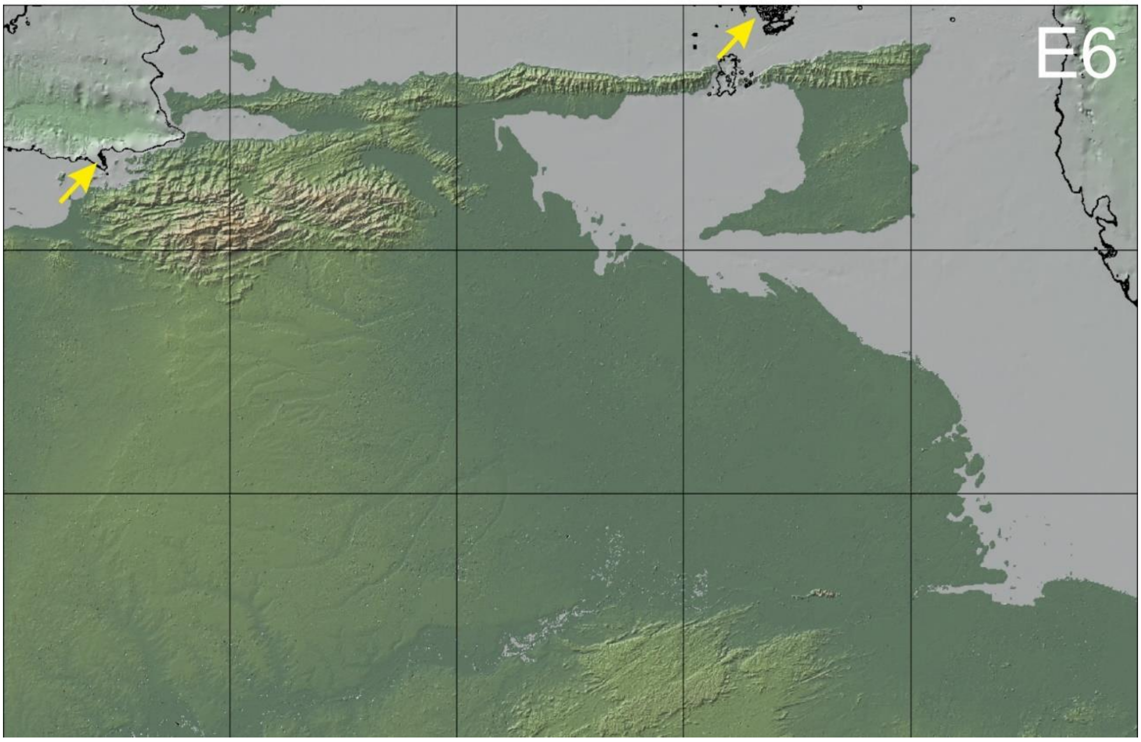
E6
